# mPFC pyramidal neuron synchrony during social competition to form social rankings is disrupted in male *Mecp2* knockout mice

**DOI:** 10.64898/2026.03.02.709145

**Authors:** Cesar Acevedo-Triana, Jennifer J. Tuscher, Jeremy J. Day, Jesus Perez-Ortega, Lucas Pozzo-Miller

## Abstract

Altered social behaviors are prevalent in neurodevelopmental disorders like monogenic Rett syndrome, which is caused by pathogenic variants in the gene encoding the methylated DNA binding transcriptional regulator MeCP2. Monosynaptic projections from the ventral hippocampus to the medial prefrontal cortex (mPFC) modulate social memory, and are altered in male *Mecp2* knockout (KO) mice. The standard tube test was used to define the social hierarchy between age- and genotype-matched triads over six consecutive days of round-robin competitions, and revealed that male *Mecp2* KO mice form social ranks but display more submissive behaviors than those observed between similarly aged triads of male wild-type (WT) littermate controls. The same triads of each genotype performed similarly in the warm spot test, where mice of each genotype compete to stand on a single warm spot in a cage with a cooled floor. The dominant WT mouse from the prior tube test had preferential and active access to the beneficial place in the competition test (warm spot) showing more dominant behaviors than the other two WT mice. On the contrary, all three *Mecp2* KO mice shared the warm spot equally, showing more submissive behaviors than those observed between the three WT mice. *In vivo* Ca^2+^ imaging from pyramidal neurons in the prelimbic mPFC during the warm spot test confirmed the presence of socially sensitive neurons, i.e., neurons that either increase or decrease their spiking activity during social interactions. mPFC pyramidal neurons in male *Mecp2* KO mice showed fewer and smaller Ca^2+^ transients during baseline, as well as during each social interaction in the warm spot test, when their activity is less synchronous than in those of WT mice. In addition, chronically inhibiting the activity of mPFC-projecting excitatory neurons of the ventral hippocampus using an intersectional DREADD approach restored behavioral deficits in male *Mecp2* KO mice. Together, these results demonstrate that male *Mecp2* mice show a low behavioral engagement during social competition tests that alters their social hierarchy and is reflected in altered activity and synchrony between mPFC pyramidal neurons. Our observations also underscore the potential relevance of this long-range projection for altered social behaviors in other mouse models of neurodevelopmental disorders associated with autism.

## Introduction

All mammals, including humans, display complex social interactions that allow individuals to function properly within groups, ensuring their survival and promoting access to resources such as food, safety, mating opportunities, and health [1, 2]. Many brain areas are associated with different forms of social behavior, including mating, aggression, and cooperation, as cooperation enables appropriate interaction with both their environment and other conspecifics [3–7]. Alterations in these brain areas can disrupt specific circuits and impair social behaviors to varying degrees. In relation to human health, altered social behaviors are common in various neuropsychiatric [8–11] and neurodevelopmental disorders (NDDs) [12–14], including those associated with autism [15, 16], where impaired sociability is a core diagnostic symptom that affects social perception and motivation, thus compromising the ability to perceive social cues and resulting in a lack of interest in others [17, 18].

In addition to deficits in social communication and language, repetitive behaviors and restricted interests are hallmark clinical features of idiopathic autism, which shows broad heterogeneity in cognitive, behavioral, genetic, and histopathological findings [19, 20]. Among NDDs with autistic features (i.e., syndromic autism), Rett syndrome (RTT) has garnered significant interest due to its association with spontaneous loss-of-function mutations in the X-linked *MECP2* gene in approximately 95% of cases [21–23]. This gene encodes the transcriptional regulator MeCP2 (methyl-CpG-binding protein 2), which regulates many critical genes involved in neuronal and synaptic function [24–27]. RTT is more prevalent in females (1:10,000) than in males, as males with complete *MECP2* dysfunction typically develop a severe encephalopathy when all brain cells lack proper MeCP2 protein function [23, 28]. Individuals with RTT develop typically for the first 6-18 months of life, followed by a period of developmental stagnation and subsequent regression of previously learned motor, language, and social skills [23, 29]. Motor impairments, debilitating stereotyped movements, breathing irregularities, sleep disturbances, and cognitive and social deficits progressively worsen in individuals with RTT [23, 24, 30]. Most of these RTT features are recapitulated in experimental mouse models based on *Mecp2* deletion and/or dysfunction, which have helped reveal the synaptic, cellular, and network bases of these behavioral deficits [31–33]. As in humans, the absence of MeCP2 in all brain cells leads to earlier onset and more severe RTT-like phenotypes in male *Mecp2* hemizygous (i.e. knockout, KO) mice compared to female heterozygous mice. Here, we followed the recommendations of the RTT/MeCP2 research community to use male *Mecp2* KO mice for fundamental studies on the consequences of *Mecp2* deletion in brain function [34].

Under laboratory conditions with unlimited water and food, adult male mice typically form a strict, stable, and transitive social hierarchy within their home cage [35, 36], and social memory is essential for identifying and distinguishing between familiar conspecifics, a crucial requirement for maintaining such stable social hierarchies [37, 38]. Using the standard linear 3-chamber test of social preference [39], we reported that symptomatic male *Mecp2* KO mice display the typical preference for a novel mouse over an inanimate object, but they exhibit impaired social memory when distinguishing between a familiar mouse and a novel mouse [40], and recently confirmed these observations in a novel 4-chamber circular arena [41]. This long-term memory of individuals within their social group depends on neuronal activity in the CA2 subregion of the dorsal hippocampus and its projections to area CA1 of the ventral hippocampus (vHIP) [42, 43], as well as vHIP projections to other regions, including the nucleus accumbens [44] and the mPFC [40]. Of relevance to RTT, mimicking the hippocampal network hyperactivity in *Mecp2* KO mice [45, 46] by chemogenetic excitation of mPFC-projecting vHIP neurons impairs social memory in WT mice, while their inhibition improves social memory in *Mecp2* KO mice [40].

Few studies have conducted in-depth analyses of social competition behaviors and their impact on social ranking in mouse models for Rett syndrome. In an early study, Shahbazian et al. (2002) reported that adult male mice with a truncated MeCP2 protein (*Mecp2^308^*) exhibited higher levels of dominant behavior, winning the majority of confrontations against wild-type (WT) mice in the tube test. Although this dominant behavior was not associated with increased aggression or enhanced social engagement during exploration in *Mecp2^308^* mice, the authors suggested two possible explanations: either WT mice actively avoided confrontation with *Mecp2^308^* mice, or *Mecp2^308^* mice were deficient in perceiving mild social threats from WT mice. Here, we show that despite their social memory deficits, male *Mecp2* KO mice can still form social hierarchies, albeit with altered dominance behaviors, which may reflect altered activity and synchrony between mPFC pyramidal neurons during social interactions, as revealed by *In vivo* Ca^2+^ imaging. In addition, chronically inhibiting the activity of mPFC-projecting excitatory neurons of the ventral hippocampus using the same intersectional DREADD approach that improved social memory in male *Mecp2* KO mice [40] also restored their hierarchal ranking deficits, underscoring the potential relevance of this long-range projection for altered social behaviors in other mouse models of neurodevelopmental disorders associated with autism.

## Results

### Mecp2 knockout mice have typical discrimination of social odors

We previously demonstrated that presymptomatic (P20) male *Mecp2* KO mice have typical sociability and social memory comparable to age-matched male WT mice [40]. After Rett-like symptom onset (P45 onward), discrimination and preference for novel over familiar mice (i.e. social memory) are impaired in the standard 3-chamber assay, and mice exhibit atypical social interactions in an unrestricted open field assay [40]. Selective chemogenetic manipulation of pyramidal neurons in the ventral hippocampus (vHIP) that project to the medial prefrontal cortex (mPFC) modulates social memory in WT mice and rescues this deficit in *Mecp2* KO mice [40]. However, the potential contribution of olfactory bulb dysfunction in *Mecp2* mutant mice [47] to this social memory deficit is unknown. We tested discrimination of social and non-social odors in *Mecp2* KO mice because odor discrimination is necessary for social identification in mice [48, 49]. We used the odor habituation/dishabituation test [50], consisting of three habituation trials with wooden blocks moistened with water (neutral odor) placed in the center of an open field, followed by a test trial with blocks impregnated with a novel odor, either chocolate (non-social) or male urine (social; Suppl. Fig. 1). *Mecp2* KO mice (N=16, P30–P40) showed discrimination of social (urine) and non-social (chocolate) odors comparable to age-matched WT littermates (N=16, P30–P40). Mixed ANOVA revealed a main effect of trial (habituation 1–3 and dishabituation) (F(3, 90) = 16.717, p < .001) and a trial × stimulus interaction (F(3, 90) = 4.588, p < .01), but no main effect of genotype or stimulus. Post hoc pairwise t-tests with Bonferroni correction showed that, for the social stimulus, both frequency of exploratory bouts (Suppl. Fig. 1B) and exploration time (Suppl. Fig. 1C) during dishabituation differed from habituation trials (habituation 2 and 3 for bout frequency; habituation 1–3 for exploration time). For the non-social stimulus, there were no differences between habituation and dishabituation trials in either genotype. There were no main effects of group (WT vs. Mecp2 KO; F(1, 30) = 2.506, p = 0.123) or stimulus type (F(1, 30) = 2.36, p = 0.13) on frequency of exploration. Similarly, there were no main effects of group (F(1, 30) = 0.947, p = 0.338) or stimulus (F(1, 30) = 2.16, p = 0.151) on exploration time. Regarding motor performance, Mecp2 KO and WT mice exhibited similar total distance traveled (t(251.63) = -1.306, p = 0.192) and speed (t(249.5) = -1.116, p = 0.265) during habituation/dishabituation. Mixed ANOVA showed no differences in distance traveled between groups (F(1, 30) = 0.322, p = 0.57), across trials (F(3, 90) = 2.25, p = 0.08), or between stimuli (F(1, 30) = 0.272, p = 0.03). Similarly, there were no main effects of group (F(1, 30) = 0.229, p = 0.635), trials (F(3, 90) = 1.02, p = 0.387), or stimulus (F(1, 30) = 1.089, p = 0.304) on speed (Suppl. Fig. 1). These findings are consistent with similar motor performance between genotypes at presymptomatic ages [51]. Together, these results indicate that social odor discrimination is preserved in Mecp2 KO mice and comparable to WT controls [40]. The absence of motor differences further supports that impaired social memory is not attributable to altered olfactory discrimination or gross motor deficits (Suppl. Fig. 1D,E).

### Mecp2 knockout mice show passive low engagement in the tube test

In a separate group of mice, social rank was assessed using the standardized tube test (i.e. competitive exclusion test), with mice housed in groups of three of the same genotype and age (P25–P40) [52–54]. After habituation and training to cross a clear tube unidirectionally, two mice were placed at opposite ends and allowed three min to compete to force the opponent out, typically yielding a winner and a loser [52, 55, 56]. Absolute social rank was determined from the percentage of wins, losses, and ties across six tournament days (Fig. 1B, 1D), classifying mice within each cage as dominant (DOM), intermediate (INT), or subordinate (SUB). In this cohort (WT = 30 [10 cages] and *Mecp2* KO = 36 [12 cages]), tournaments were conducted over six consecutive days among cage mates. WT-DOM mice won 81.25% of trials and lost 15.42%; WT-INT mice won 51.67%, lost 44.17%, and tied 4.17%; WT-SUB mice won <12%, lost 80.42%, and tied 8.33% (Fig. 1B). *Mecp2* KO-DOM mice won 61%, lost 6%, and tied >33%; *Mecp2* KO-INT won 36%, lost 29%, and tied 35%; *Mecp2* KO-SUB won 10%, lost 70%, and tied 20%. Hierarchical distance was estimated using EloRating scores. Mixed ANOVA showed a main effect of rank (F(2, 60) = 112.7, p < .001) and a rank × day interaction (F(10, 300) = 34.27, p < .001), with no main effect of genotype (F(1, 60) = 0.001, p = .978) or day (F(5, 300) = 0.005, p = .999). Post hoc pairwise t-tests with Bonferroni correction indicated separation of DOM, INT, and SUB ranks in both genotypes by day three (Fig. 1C, 1D). *Mecp2* KO mice exhibited a higher percentage of ties across ranks.

**Figure 1.**
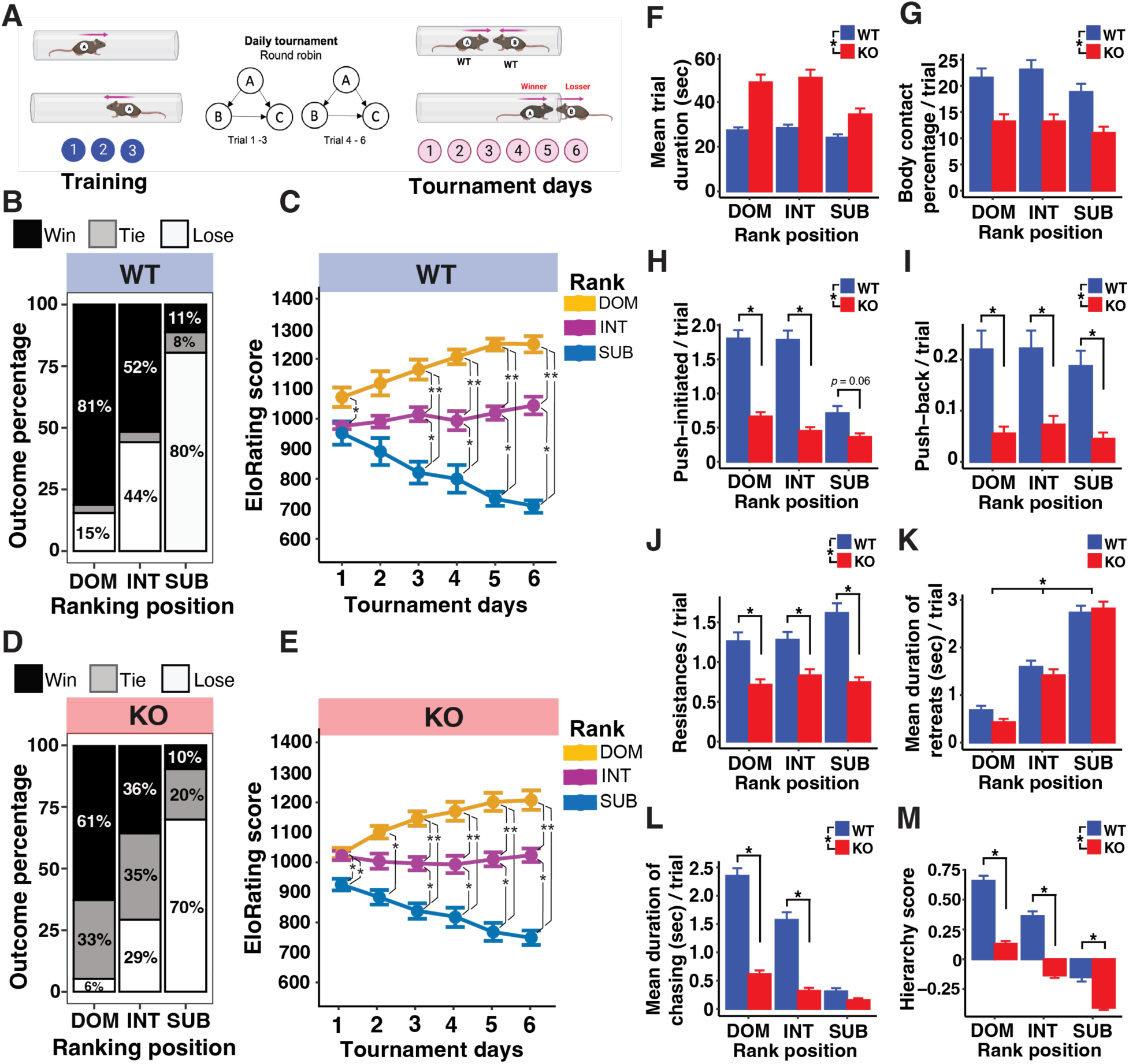
Male *Mecp2* KO mice form social ranks with lower hierarchy score and showing more submissive behaviors among them. (A) Social hierarchy between 3 age-matched male mice of the same genotype was determined using the tube test (3 days of training alone and 6 days of round-robin style tournaments between male mice of the same genotype)5 (Fig 2A). (B, D) Cumulative percentages of trial outcomes classifying the 3 male WT mice (B) and male *Mecp2* KO mice (D) in each cage into dominant (DOM), intermediate (INT), and subordinate (SUB), based in their percentages of wins, loses, and ties after 6 days of tournament. (C, E) EloRating score in each tournament day for male WT mice (C) and male *Mecp2* KO mice (E); EloRating scores start as 1,000. Analyzing performance during the tube test tournament, we found that male *Mecp2* KO mice have atypical behaviors during the tube test to solve this social conflict: they spent more time inside the tube (F), engaged in fewer body contacts inside the tube (G), initiated fewer *pushes* (H), and fewer *push-back* as response to pushing (I), engaged in fewer *resistances* (J), no differences during the *retreat* time (K), and spent less time *chasing* the opponent (L). Hierarchy score was calculated as a composed measurement of dominant behaviors, similar to previously reported [67, 164]. We calculated the z-scores from total number of wins during pairwise tournament, *pushing*, *push-back*, *resistance* and *chasing* behavior. Once each z-score were calculated were combined in the hierarchy score. A high punctuation in this hierarchy score means that mice showed more dominant behaviors after tournament, punctuations close to zero or negative mean submissive or lack of dominance. Data in F-M were analyzed with a mixed factorial ANOVA model. * p < 0.05

To further characterize the performance in the tube test, several behaviors were scored [52, 53, 57], including *push-initiated*, *push-back*, *resistance*, *chasing*, and *retreat* (Fig. 1). *Trial duration* showed a main effect of genotype (F(1, 60) = 5.49, p = .002), with no effect of rank (F(2, 60) = 0.711, p = .494) or genotype × rank interaction (F(2, 60) = 0.238, p = .788) (Fig. 1F). WT mice resolved trials in less time than *Mecp2* KO mice, independent of rank. *Body contact* showed a main effect of genotype (F(1, 60) = 5.456, p = .022), with no effect of rank (F(2, 60) = 0.269, p = .765) or interaction (F(2, 60) = 0.024, p = .975) (Fig. 1G). WT mice spent more time in contact with opponents than *Mecp2* KO mice. *Push-initiated* behavior showed main effects of genotype (F(1, 60) = 164.46, p < .001) and rank (F(2, 60) = 30.64, p < .001), and a genotype × rank interaction (F(2, 60) = 16.91, p < .001) (Fig. 1H). Tukey’s HSD tests showed differences between WT-DOM vs. *Mecp2* KO-DOM (diff = -1.131; p < .0001) and WT-INT vs. *Mecp2* KO-INT (diff = -1.133; p < .0001), but not WT-SUB vs. *Mecp2* KO-SUB (diff = -0.347; p = .068). *Push-back* behavior showed a main effect of genotype (F(1, 60) = 57.588, p < .001), with no effect of rank (F(2, 60) = 0.821, p = .044) or interaction (F(2, 60) = 0.107, p = .899). Tukey’s HSD indicated differences between WT and *Mecp2* KO within each rank (WT-DOM vs. *Mecp2* KO-DOM diff = -0.164; p < .001; WT-INT vs. *Mecp2* KO-INT diff = -0.148; p < .001; WT-SUB vs. *Mecp2* KO-SUB diff = -0.142; p < .001). *Resistance during pushing* showed a main effect of genotype (F(1, 60) = 72.586, p < .001), no main effect of rank (F(2, 60) = 2.031, p = .132), and a genotype × rank interaction (F(2, 60) = 3.064, p = .047) (Fig. 1J). Tukey’s HSD showed differences between WT and *Mecp2* KO within each rank (WT-DOM vs. *Mecp2* KO-DOM diff = - 0.546; p < .001; WT-INT vs. *Mecp2* KO-INT diff = -0.447; p < .01; WT-SUB vs. *Mecp2* KO-SUB diff = -0.87; p < .001). *Retreat duration* showed a main effect of rank (F(2, 60) = 170.496, p < .001), with no effect of genotype (F(1, 60) = 1.277, p = .259) or interaction (F(2, 60) = 1.059, p = .347) (Fig. 1K). *Chasing duration* showed main effects of rank (F(2, 60) = 96.42, p < .001) and genotype (F(1, 60) = 227.36, p < .001), and a genotype × rank interaction (F(2, 60) = 45.07, p < .001) (Fig. 1L). A grouped hierarchical score was calculated using z-scores of wins, push-initiated, push-back, resistance, retreat, and chasing behaviors. Mixed ANOVA showed main effects of genotype (F(1, 1576) = 311.59, p < .001) and rank (F(1, 1576) = 252.40, p < .001), and a genotype × rank interaction (F(2, 1576) = 12.54, p < .001) (Fig. 1M).

Taken together, these ethogram analyses clearly show that dominant behaviors (*pushing*, *resistance*, *chasing*) followed a hierarchical pattern in WT mice (DOM > INT > SUB), while *Mecp2* KO mice exhibited fewer dominant behaviors and lower hierarchical scores than their age-matched WT controls.

### Mecp2 KO mice do not use social rank information to solve access to a preferential place

To confirm the passive hierarchy profile observed during social rank formation in *Mecp2* KO mice using the tube test, we next used the warm spot test, in which mice are placed in an open arena with a cold floor and compete for access to a warm spot that fits only one mouse at a time [52–54]. The same group of three WT and three *Mecp2* KO mice that had completed the tube test were tested 24 h after the final tournament day (Fig. 3A–C). Previous studies show that higher-ranked mice spend more time on the warm spot than lower-ranked mice [52, 54, 58], consistent with social rank influencing access to resources [54, 59]. WT mice accessed the warm spot more frequently than *Mecp2* KO mice. Mixed ANOVA showed a main effect of genotype (F(1, 60) = 6.427, p = .013) (Fig. 3D), with no effect of rank (F(2, 60) = 1.615, p = .276) or genotype × rank interaction (F(2, 60) = 1.792, p = .175). Tukey’s HSD tests showed no differences within ranks (WT-DOM vs. *Mecp2* KO-DOM diff = −3.572; p = .055; WT-INT vs. *Mecp2* KO-INT diff = −1.483; p = .820; WT-SUB vs. *Mecp2* KO-SUB diff = 1.522; p = .786). WT mice also spent more time on the warm spot. Mixed ANOVA showed main effects of genotype (F(1, 60) = 5.784, p = .019) and rank (F(2, 60) = 5.954, p = .004), with no genotype × rank interaction (F(2, 60) = 1.518, p = .227) (Fig. 3E). Tukey’s HSD tests showed no pairwise differences. Correlations between wins in the tube test and warm spot duration were low and not significant in either WT (rho = .269, p = .149) or *Mecp2* KO mice (rho = .119, p = .486) (Fig. 3F).

**Figure 2.**
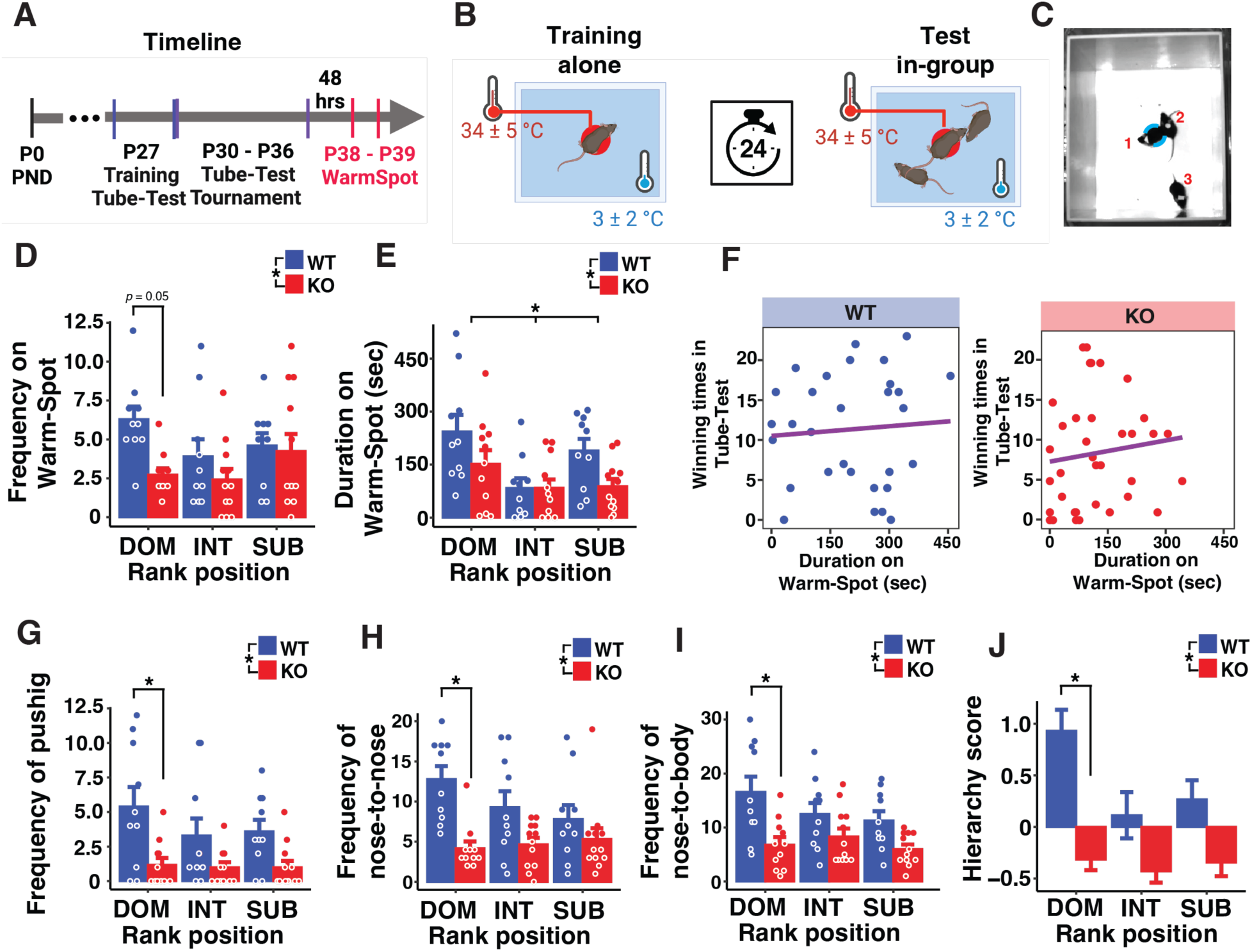
Male *Mecp2* KO mice showed fewer frequency and low duration on the warm spot, and low hierarchy score during the warm spot test. (A) To confirm whether social ranking is used to solve social conflicts, the warm spot test was performed during 10 min, previously mice were individually trained in the same warm spot box to sure that each one know where the warm spot was located (B), 48 h posterior to the last day of the tube test tournament. (D, E) In contrast to male WT mice where the DOM mouse stayed the longest on the warm spot (D) and for more instances in each trial (E), male *Mecp2* KO mice spent similar amounts of time on the warm spot and for fewer instances, irrespective of their social rank (previously determined in the tube test). (F) The number of winning trials in tube test shows a similar positive correlation with the time spent on the warm spot but without differences between groups. (G - I) During social interactions in the warm spot test *Mecp2* mice showed fewer *pushing* (G), fewer *nose-to-nose* (H), and fewer *nose-to-body* interactions (I). (J) Equivalent to the hierarchy score during the tube test, the composed measurement of dominant behaviors during the warm spot test was calculated with z-score from frequency and duration on the warm spot, frequency of *pushing*, frequency of *nose-to-nose*, and frequency of *nose-to-body*. Panels D, E, G, H, I and J show a main effect of group, but not of social rank except to duration on warm spot. * p < 0.05.

**Figure 3.**
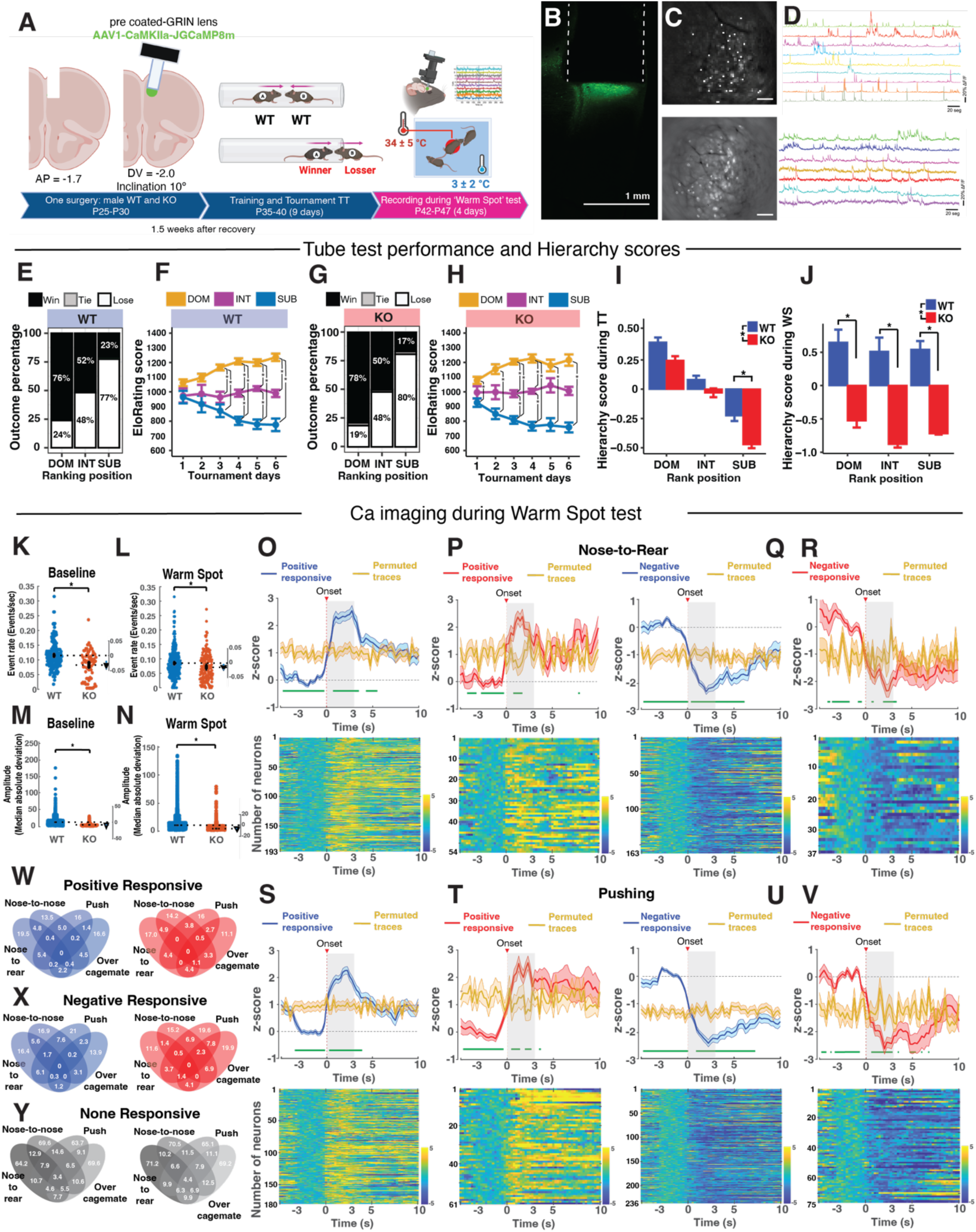
Pyramidal mPFC neurons in male *Mecp2* KO mice showed fewer and smaller Ca^2+^ transients during baseline, as well as during social interactions in the warm spot test, but a similar proportion of social-sensitive neurons. (A) Diagram of modified surgery to implant pre-coated-GRIN lens in the mPFC of WT and *Mecp2* KO mice, followed by tube test tournament to determinate their social ranking, and the warm spot test 24 hours after. (B) Example of pre-coated-GRIN lens placement and jGCaMP8m-expressing excitatory neurons (*CaMKIIa* promoter). (C) Representative FOV of maximal projection in WT mouse (C top) after motion correction and CNMFe processing. (D top) Representative jGCaMP8m traces (% dF/F) during the training in the warm spot test, neurons were randomly selected in each genotype. (C bottom) Representative FOV of maximal projection of *Mecp2* KO mice. (D bottom) Representative jGCaMP8m traces (% dF/F) of *Mecp2* KO mice. (E - H) Cumulative percentages of trial outcomes classifying the 3 male WT mice (E) and male *Mecp2* KO mice (G) in each cage into dominant (DOM), intermediate (INT), and subordinate (SUB). (F, H) EloRating score in each tournament day for male WT mice (F) and male *Mecp2* KO mice (H); (I, J) Hierarchy scores calculated during the tube test (I), and after the warm spot test (J) showed that WT mice exhibited more dominant behaviors after tournament and during the warm spot test. (K, L) The frequency of jGCaMP8m events during habituation (baseline) and the entire warm spot test was significantly lower in *Mecp2* KO mice than in WT mice. (M) The amplitude of jGCaMP8m events was significantly lower in *Mecp2* KO mice during both habituation and the warm spot test. Behavioral movies were annotated and synchronized with jGCaMP8m movies using BENTO software [63]. To detect neurons responsive to each behavior, each signal above or below two standard deviations from the z-score normalized using a baseline of 3 seconds prior to the onset of each behavior was considered as positively or negatively responsive, respectively. The average trace was compared with the same trace permuted, with the green bar below traces denoting significance p-values calculated by false discovery rates. False Discovery Rates (FDR) compared point-to-point t-tests and corrected for multiple comparisons using significant values satisfying P(k) < 0.05 * k/2000. (O, P) Average dF/F z-scores during nose-to-rear interaction showing average of traces classified as positive responsive in WT mice (O) and in *Mecp2* KO (P). (O bottom) Heatmap of dF/F z-scores during nose-to-rear exploration in WT mice, (P bottom) in *Mecp2* KO mice. (Q, R) average dF/F z-scores during nose-to-rear interaction showing average of traces classified as negative responsive in WT mice (Q top) and in *Mecp2* KO (R). (S, T) Positive responsive traces in WT mice (S top) and in *Mecp2* KO mice (T top) during *pushing* behavior. Negative responsive (U and V). (W -Y). Percentage of positive, negative and non-responsive neurons during different social behaviors was similar in the mPFC of *Mecp2* KO (n = 9/21 mice) and WT mice (n = 13/21 mice). Scale bar in panel C correspond to 100 μm. * p < 0.05.

Dominant and social behaviors during the warm spot test were also scored. *Pushing* was more frequent in WT than *Mecp2* KO mice. Mixed ANOVA showed a main effect of genotype (F(1, 60) = 19.161, p < .001) (Fig. 3G), with no effect of rank (F(2, 60) = 1.083, p = .345) or interaction (F(2, 60) = .734, p = .484). Tukey’s HSD showed a difference for WT-DOM vs. *Mecp2* KO-DOM (diff = −4.218; p = .011), but not for INT or SUB comparisons. *Nose-to-nose exploration* showed a main effect of genotype (F(1, 60) = 20.170, p < .001) (Fig. 3H), with no effect of rank (F(2, 60) = 0.946, p = .394) or interaction (F(2, 60) = 2.401, p = .099). Tukey’s HSD showed a difference for WT-DOM vs. *Mecp2* KO-DOM (diff = −8.618; p = .001), but not for other ranks. *Nose-to-body exploration* showed a main effect of genotype (F(1, 60) = 19.736, p < .001) (Fig. 3I), with no effect of rank (F(2, 60) = 1.544, p = .222) or interaction (F(2, 60) = 1.411, p = .252). Tukey’s HSD showed a difference for WT-DOM vs. *Mecp2* KO-DOM (diff = −9.781; p = .003), but not for INT or SUB comparisons.

Additional measures were consistent with reduced social engagement in *Mecp2* KO mice. *Resting* duration showed a main effect of genotype (F(1, 60) = 8.286, p = .005), with no effect of rank (F(2, 60) = 0.090, p = .913) or interaction (F(2, 60) = 1.923, p = .155). *Nose-to-rear exploration* showed main effects of genotype (F(1, 60) = 12.510, p < .001) and rank (F(2, 60) = 3.317, p = .043), with no interaction (F(2, 60) = 0.205, p = .814). *Grooming* showed a main effect of genotype (F(1, 60) = 8.799, p = .004), with no effect of rank (F(2, 60) = 1.454, p = .241) or interaction (F(2, 60) = 1.276, p = .286). A grouped hierarchical score for the warm spot test was calculated using z-scores of warm spot frequency and duration, *pushing*, *nose-to-nose*, and *nose-to-body* exploration. Mixed ANOVA showed main effects of genotype (F(1, 60) = 37.740, p < .001) and rank (F(2, 60) = 3.976, p = .023), with no genotype × rank interaction (F(2, 60) = 2.973, p = .058) (Fig. 3J).

Overall, WT mice displayed higher frequencies and durations on the warm spot and more dominant and social exploratory behaviors than *Mecp2* KO mice. *Mecp2* KO mice showed lower hierarchical scores and reduced social engagement. These results parallel those from the tube test and indicate reduced participation in social competition in *Mecp2* KO mice.

### Pyramidal mPFC neurons in Mecp2 KO mice are less active but the proportions of social-sensitive neurons are similar to WT mice

Previous studies have described that some neurons in the mPFC are preferentially activated or inhibited during social interactions [60]. In agreement with these findings, we identified groups of mPFC excitatory neurons that were active during social interactions in WT mice. In these pilot experiments, P21–P25 male WT (n = 3) mice received unilateral mPFC injections of AAVs expressing the Ca²⁺ indicator jGCaMP6s under control of the *CamkIIa* promoter; a GRIN lens/baseplate assembly was implanted over the injection site in the mPFC one week later (Suppl. Fig. 3). We confirmed the presence of socially sensitive neurons in the prelimbic (PL) area of the mPFC that were either activated (social-ON neurons) or inhibited (social-OFF neurons) during close exploration of a novel mouse, as previously reported [60]. Examples of excitatory mPFC neurons that increased the frequency of spontaneous Ca²⁺ transients during a 2-second time window before and after close exploration of a novel mouse in a novel 4-chamber arena are shown in Suppl. Fig. 3F (bottom red traces). These neurons exhibited activity rates approximately three times higher than during non-social behaviors. Examples of mPFC excitatory neurons that decreased the frequency of spontaneous Ca²⁺ transients during social exploration are also shown in Suppl. Fig. 3F (top blue traces).

After confirming the involvement of the mPFC in social interactions in WT mice, we aimed to determine whether these socially sensitive neurons were altered in *Mecp2* KO mice. To reduce the number of surgeries in all mice, especially *Mecp2* KOs, we pre-coated GRIN lenses with a mixture of silk fibroin [61] and AAV-expressing jGCaMP8m (*CamkIIa* promoter) [62] before their implantation into the mPFC of WT (n = 21) and *Mecp2* KO (n = 21) mice (Suppl. Fig. 3A and Suppl. Fig. 4A–D). Approximately 10 days after surgery, both WT and *Mecp2* KO mice were habituated and trained in the tube test, followed by a 6-day pairwise round-robin competition among three age-matched mice (maximum 3 minutes per trial), using the same protocol as in previous experiments. The cumulative win percentage and daily EloRating scores indicated that the surgical procedures and implanted GRIN lens did not affect ranking formation (Fig. 3E–H). The percentages of wins, ties, and losses were similar between WT and *Mecp2* KO mice, suggesting preserved ranking formation. Of note, *Mecp2* KO mice in this cohort showed a lower rate of ties in the tube test compared to the previous cohorts of *Mecp2* KO mice without miniscopes, during which ties were commonly observed between them. We suspect the added size of the miniscope headpiece limited the negotiation in the tube test, and allowed passing-by outcomes (i.e. ties) less feasible and common.

Regarding daily performance during the tube test that determines the EloRating score (Fig. 3F and H), a mixed ANOVA showed a main effect of rank (F (2, 36) = 6.106, p = .056), but no main effect of genotype (F (1, 36) = 0.874, p = .356), day of tournament (F (5, 180) = 0.969, p = .437), or interactions between genotype, rank, and day. Post hoc comparisons by pairwise t-test with Bonferroni correction of EloRating scores showed that both WT and *Mecp2* KO mice exhibited differences between DOM, INT, and SUB mice after the third day of tournament, similar to the previous cohort of mice without miniscopes. The grouped hierarchical score after the tube test replicated previous findings, showing a main effect of genotype (F (1, 1002) = 14.986, p < .001) (Fig. 3I), a main effect of ranking (F (1, 1002) = 90.124, p < .001), and a significant interaction between genotype and rank (F (2, 1002) = 12.54, p < .001). Post hoc comparisons using Tukey’s HSD test showed differences between WT-SUB vs *Mecp2* KO-SUB (diff = −.171; p = .001), but no differences between WT-INT vs *Mecp2* KO-INT (diff = −0.04; p = .956) or WT-DOM vs *Mecp2* KO-DOM (diff = −0.130; p = .109). In contrast, the hierarchical score after the warm spot test showed a main effect of genotype (F (1, 19) = 99.307, p < .001), but no main effect of rank (F (1, 19) = 1.314, p = .292) or interaction between genotype and rank. Post hoc comparisons using Tukey’s HSD test showed differences between WT-DOM vs *Mecp2* KO-DOM (diff = −1.175; p < .001), WT-INT vs *Mecp2* KO-INT (diff = −1.391; p < .001), and WT-SUB vs *Mecp2* KO-SUB (diff = −1.261; p = .002). These data confirm that *Mecp2* KO mice display lower hierarchical scores, more subordinate behaviors, and reduced social exploration and engagement with their genotype-matched *Mecp2* KO cagemates. For the most part, these traits were not altered by surgical implantation for *in vivo* Ca^2+^ imaging with a head-mounted miniscope, except the lower rate of ties between *Mecp2* KO mice when carrying the miniscope headpiece.

During the warm spot test, mouse behavior was recorded with an IR-sensitive sCMOS camera and synchronized with acquisition of jGCaMP8m movies with the head-mounted miniscope. The miniscope imaging software (IDEAS, Inscopix, CA) was synchronized with behavioral videos acquired using a top-view sCMOS camera controlled by EthoVision (Noldus) via TTL pulses. Individual jGCaMP8m events were detected using a threshold of 4 median absolute deviations (MAD) from the original trace, assuming a minimum exponential decay time of 0.20 s. A total of 1,216 neurons were imaged from 13 WT mice (21 mice/7 cages) and 389 neurons were imaged from 9 *Mecp2* KO mice (21 mice/7 cages). The frequency of these jGCaMP8m events during habituation (baseline) (t-test = 5.06, p < .001) (Fig. 3K) and during the entire warm spot test (t-test = 2.99, p = .002) (Fig. 3L) was lower in *Mecp2* KO mice than in WT mice. In addition, the amplitude of individual jGCaMP8m events was smaller in *Mecp2* KO mice during both habituation (t-test = 28.72, p < .001) and the warm spot test (t-test = 48.15, p < .001). Behavioral videos were synchronized with miniscope Ca²⁺ movies using the *Behavior Ensemble and Neural Trajectory Observatory* (BENTO) software [63], enabling synchronized behavioral annotation across seven recorded behaviors. These behaviors were consistent with those scored using BORIS in the previous cohorts, including the frequency and duration on the warm spot, social exploration (nose-to-nose, nose-to-rear, nose-to-body), pushing, climbing over a cagemate, chasing/following, grooming, and jumping.

To identify neurons participating in specific behaviors, raw CNMFe traces were aligned to each behavior, averaged per neuron, and normalized using Z-scores relative to each neuron’s baseline activity (3 seconds prior to behavior onset). Neurons were classified as increasing or decreasing their activity using a threshold of 2 standard deviations (95% confidence interval of a normal distribution) based on the Z-score within 3 seconds after behavior onset. This threshold was used to classify neurons as positively responsive, negatively responsive, or non-responsive [64], consistent with previous social behavior studies [65, 66]. To verify classification accuracy, the average jGCaMP8m ΔF/F trace identified as positive or negative was compared with a permutated version of the same ΔF/F trace. Significant p-values were calculated using false discovery rate (FDR) correction (Fig. 3O–V, top). FDR correction was applied to point-to-point t-tests for multiple comparisons using the criterion P(k) < 0.05 × k/2000 [67]. Overall, there were no differences in the proportion of neurons classified as positive, negative, or non-responsive compared to permutated traces (Fig. 3O–V; Suppl. Fig. 4E–F) between either WT or *Mecp2* KO mice. Given the different total numbers of neurons analyzed (WT: n = 1,216; *Mecp2* KO: n = 389), percentages were used for comparisons. During nose-to-rear interaction, the percentage of positively responsive neurons in WT mice (19.5%, n = 193) was similar to that in *Mecp2* KO mice (17.0%, n = 54). Similar results were observed for negatively responsive neurons (WT: 16.4%, n = 163; *Mecp2* KO: 11.6%, n = 37). Although WT mice showed greater engagement during social competition in the warm spot test than *Mecp2* KO mice, the percentage of neurons classified as positively responsive during *pushing* behavior was similar between WT (16%, n = 180) and *Mecp2* KO (16%, n = 61) mice. The percentage of negatively responsive neurons during *pushing* was higher than that of positively responsive neurons, but was also similar between WT (21%, n = 236) and *Mecp2* KO (19.6%, n = 75) mice.

These findings were consistent across all behaviors analyzed. Pearson’s chi-squared tests showed no differences in the percentage of responsive neurons between genotypes for *nose-to-rear* (χ²(2) = 1.3603, p = .506), *nose-to-body* (χ²(2) = 1.026, p = .598), *nose-to-nose* (χ²(2) = .109, p = .946), *pushing* (χ²(2) = .055, p = .972), or *climbing over a cagemate* (χ²(2) = 2.128, p = .345) (Fig. 3W–Y). Similar results were obtained when neurons were classified according to social rank (Suppl. Fig. 4G–H). These results indicate that, although dominant behaviors were more frequent in WT mice, this was not associated with a higher proportion of behaviorally responsive mPFC excitatory neurons. The relatively low percentage of socially sensitive neurons is consistent with previous reports showing that only approximately 10% of excitatory mPFC neurons are socially responsive [60, 68].

### Functional co-activation of mPFC pyramidal neuron ensembles during social interactions is lower in Mecp2 KO than in WT controls

To determine whether mPFC pyramidal neurons identified as social-sensitive exhibited similar patterns of correlated activity during social interactions, we conducted a correlation analysis for each mouse and compared it across social behaviors. Pilot correlations revealed that groups of neurons were correlated with one another, forming clusters of similar activation patterns (Suppl. Fig. 5). Because neuronal ensembles have been proposed as groups of neurons that fire together and represent neural encoding for memory, perception, motor function, and behavior [69–71], we investigated how many neurons could be co-activated within the same ensemble, and whether such co-activation could sufficiently represent the social interactions observed during the warm spot test. Individual Ca^2+^ transients were inferred from jGCaMP8m ΔF/F traces (CNMFe traces) using deconvolution with a threshold of three SNR, applying the OASIS algorithm (*Online Active Set Method to Infer Spikes*) [57] to produce binarized signals (Fig. 4A). Following Perez-Ortega et al. [70, 72], ensembles were identified based on functional connectivity between pairs of neurons that were co-activated within a 10 ms time window, surpassing the threshold of 1,000 spike raster surrogates generated by random circularization of active frames. Each significant co-activation frame (one frame bin = 10 ms) was considered a vector. The similarity between each pair of vectors was assessed using the *Jaccard* similarity index, where a value near zero indicated entirely different neuronal composition between vectors and a value of one indicated identical composition [70, 72]. The activity patterns of these vectors were clustered by hierarchical clustering, grouping co-activated frames with similar patterns while discarding clusters falling significantly below raster surrogates (Fig. 4G and 4K). These clusters were interpreted as ensembles encoding functional co-activation of frames.

**Figure 4.**
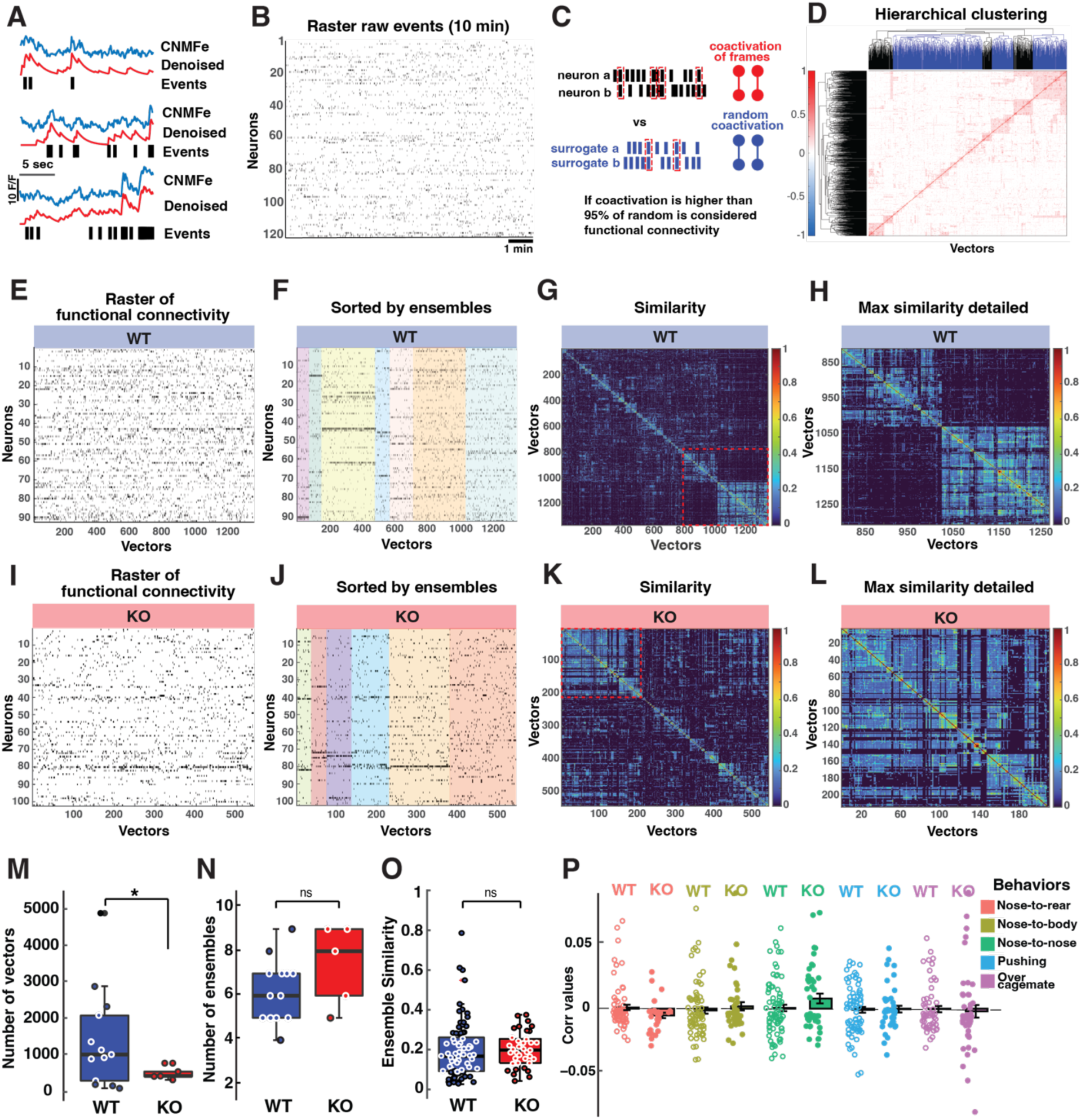
Male *Mecp2* KO mice showed fewer functional coactivations during the warm spot test. (A) Diagram of events/spikes inferred from deconvolution processes to obtain a binarized signal from each neuron. (B) Representative raster plot showing events across the entire session (10 mins). (C) Method used to identify functional connectivity based on the coactivation of frames between pairs of neurons. The activity of each pair of neurons was compared with 1000 surrogates by a circular shifting their activity (Perez-Ortega, 2021, 2024). A cumulative distribution probability of 95% from the surrogate coactivation was then compared to real coactivation; if real coactivation exceeded this threshold, it was considered significant. (D) Hierarchical clustering of pairwise co-activations (vectors) grouped by similar coactivation patterns. (E and I) Representative WT and KO examples of filtered raster based on functional connectivity. (F and J) Rasters of functional connectivity sorted by ensembles, identified by color. (G and K) Column vectors from raster sorted by hierarchical clustering using single linkage based on Jaccard similarity. (H and L) Max similarity showed in G and K, respectively, indicating groups of vectors with high levels of similarity. (M) WT mice showed a significantly higher number of pairwise vectors considered to be functionally coactivated across all ensembles. (N) No differences were observed between WT and KO mice in the number of ensembles per session. (O) Ensemble similarity represents the average Jaccard similarity between each pairwise vector of the ensemble raster matrix. No differences were observed in ensembles similarity between WT and KO mice. (P) Correlation between ensembles and each social behavior during the ‘Warm Spot’ test.

The number of vectors (i.e., functional co-activations) during the entire warm spot test was significantly lower in *Mecp2* KO mice (M = 490.8 vectors) than in WT mice (M = 1,399.07 vectors; Welch t-test = 2.345, p = .035; Fig. 4M), indicating impaired functional co-activation within mPFC excitatory neuron ensembles in *Mecp2* KO mice. However, neither the number of ensembles (Welch t-test = −1.294, p = .243; Fig. 4N) nor the similarity within ensembles (Welch t-test = 0.315, p = .752; Fig. 4O) differed significantly between genotypes, suggesting that the reduction in functional co-activation between mPFC excitatory neurons in *Mecp2* KO mice was not attributable to fewer ensembles or lower intra-ensemble coherence.

To determine whether different social behaviors could be encoded by specific ensembles, we compared the correlation between behavior frequency and ensemble activity during those behaviors. Correlation coefficients were low and highly variable (Fig. 4P), indicating low specificity of mPFC excitatory pyramidal neuron ensembles in the prelimbic area for encoding individual social behaviors, with no significant difference between genotypes.

These findings indicate that *Mecp2* KO mice exhibit lower functional co-activation within mPFC pyramidal neuron ensembles during the warm spot test compared to WT controls, despite equivalent ensemble numbers and within-ensemble similarity across genotypes. Moreover, functional co-activation within mPFC pyramidal neuron ensembles does not appear to encode specific social behaviors, in contrast to findings in primary sensory areas such as visual cortex [70, 72] and auditory cortex [73]. This lack of specificity may reflect the complex convergent inputs the mPFC receives from multiple brain regions [74, 75]. Together, these observations reinforce the view of the mPFC as an integrative associational cortex that processes multiple emotional and cognitive inputs [15, 40, 60, 76–78], and underscore the behavioral relevance of mPFC dysfunction in *Mecp2* KO mice.

### Ensembles of mPFC pyramidal neurons do not encode the odor identity of cagemates and remain stable during the formation social ranking in both Mecp2 KO and WT mice

To test whether the functional co-activations observed during the warm spot test reflected the higher frequency of social behaviors in WT mice rather than stimulus-specific encoding, and to determine whether mPFC excitatory neurons could encode specific social stimuli, we exposed a separate cohort of mice (WT = 3; *Mecp2* KO = 2) to the urine odor of their cagemates during calcium imaging in a head-fixed preparation. The number of odor exposures was controlled across the last five days of the tube test tournament (days 2, 3, 4, 5, and 6; Fig. 5A). Each mouse was randomly exposed ten times to each of three odors: cagemate urine odor A, cagemate urine odor B, and a control vanilla odor. The first two minutes served as habituation, with odor exposure beginning two minutes after placement in the head-fixed device. Each trial consisted of a 2-s baseline, followed by 2 s of odor exposure and 18 s of post-exposure recording [79]. Individual Ca^2+^ transients were inferred from CNMFe traces (jGCaMP8m ΔF/F) using the OASIS algorithm [57] to generate binarized signals, and ensembles were identified from functional co-activations following Perez-Ortega et al. (2021, 2024) [70, 72].

**Figure 5.**
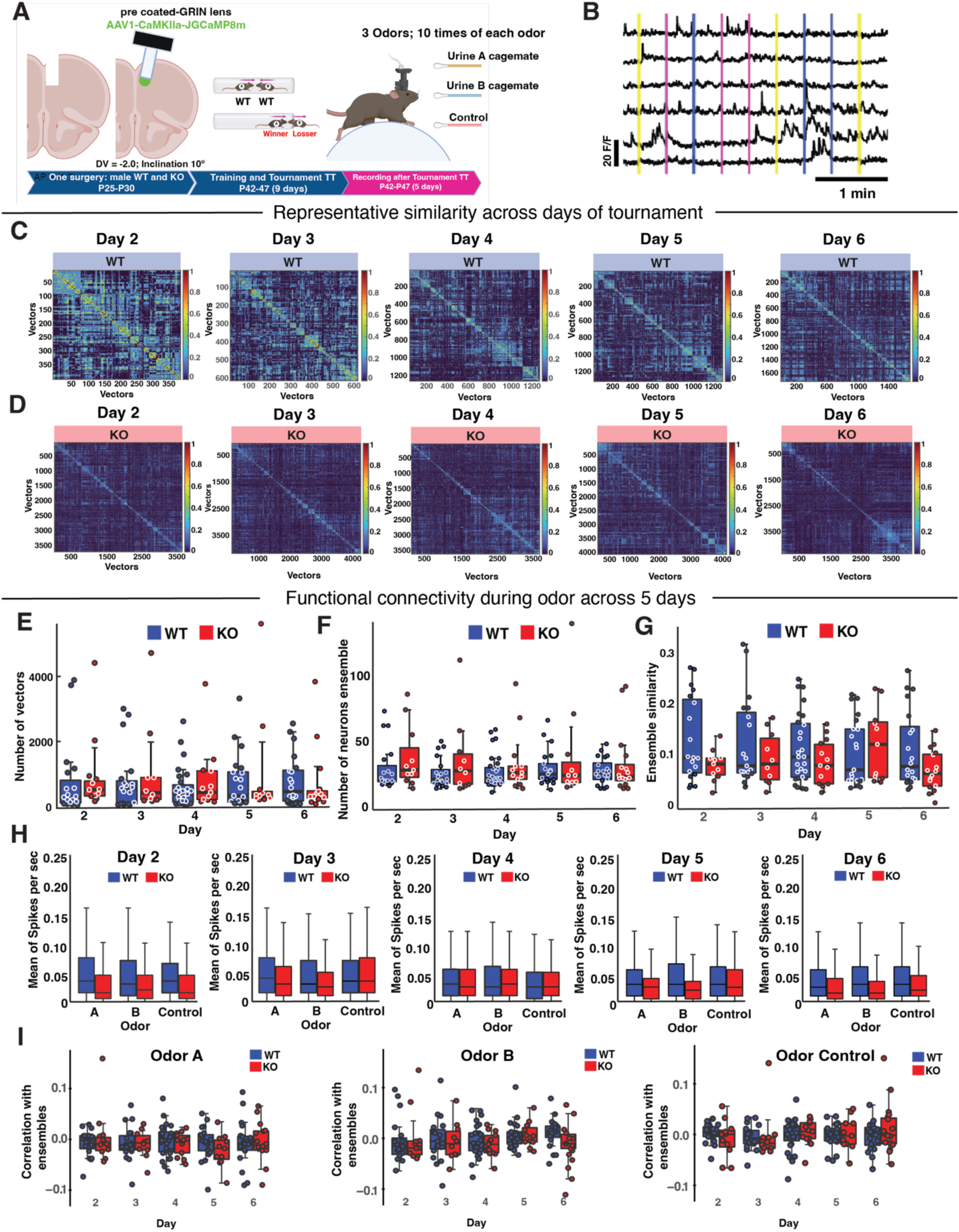
Ensembles of pyramidal mPFC neurons do not encode urine odor of cagemates and remain stable during social ranking formation. (A) Diagram of surgery to implant pre-coated GRIN lens in the mPFC of WT and *Mecp2* KO mice, followed by posterior tube test tournament to determine the social ranking. One hour after TT tournament mice were imaging (day 2 to day 6 of TT tournament) while they were exposed to the urine odor of cagemates while they were head-fixed mice. Each mouse was exposed to two cagemates’ urine odors and control odor 10 times per session. Each odor was delivered for 2s followed by 18 s without odor in a randomized trial each day. (B) Representative random calcium traces with the overlaid timing of odor presentations. (C) Representative similarity results across day 2 to day 6 of imaging of the same WT mouse. (D) Representative results of similarity in a *Mecp2* KO mouse. (E) Number of vectors across every day. (F) Number of neurons that participate in ensembles every day. (G) Similarity within ensembles. (H) The mean of spikes per second was inferred from the binarization of the raster plot. (I) Correlation of ensembles during each odor exposition.

We hypothesized that WT mice would exhibit greater within-ensemble similarity and a higher number of vectors across days, given their more pronounced hierarchical profile during social competition observed in both the tube test tournament and the warm spot test. This hypothesis was motivated by two lines of evidence. First, hierarchical social behaviors such as *pushing*, *resistance*, and *chasing* are associated with increased mPFC activity [54, 59, 80] and with changes in the physiological properties of mPFC pyramidal neurons [53, 81]. Second, mPFC pyramidal neurons have been shown to encode information about social opponents during tube test competition [82, 83]. Furthermore, our earlier results indicated that odor discrimination is preserved in *Mecp2* KO mice, and that functional co-activation during the warm spot test was greater in WT than in *Mecp2* KO mice. Contrary to this hypothesis, a mixed ANOVA revealed no significant main effect of genotype (F(1, 3) = 0.127, p = .744), day (F(4, 12) = 1.238, p = .346), or their interaction (F(4, 12) = 0.965, p = .461) on the number of vectors during odor exposure (Fig. 5E), indicating that WT and *Mecp2* KO mice exhibited similar co-activation patterns across all days. Similarly, no significant effects were found for the number of ensembles: genotype (F(1, 3) = 2.056, p = .247), day (F(4, 12) = 0.516, p = .725), or their interaction (F(4, 12) = 0.348, p = .815; Fig. 5F). In contrast, within-ensemble similarity showed a significant genotype × day interaction (F(4, 12) = 4.026, p = .026; Fig. 5G), with no significant main effects of genotype (F(1, 3) = 0.214, p = .674) or day (F(4, 12) = 1.791, p = .195), suggesting that ensemble similarity changes across tournament days in a genotype-dependent manner.

To determine whether pyramidal neurons differentially encoded urine odors versus control across tournament days, a mixed ANOVA revealed a significant genotype × odor interaction (F(2, 14436) = 4.591, p = .01), a significant main effect of day (F(4, 12) = 4.361, p = .02), and a significant odor × day interaction (F(8, 14436) = 2.857, p = .003; Fig. 5H), indicating that odor encoding by mPFC pyramidal neurons is both genotype- and day-dependent. However, correlation analysis across odor exposures and days revealed no significant differences in correlation coefficients (Fig. 5I), consistent with low ensemble specificity for individual odor identity. Taken together, these results indicate that functional co-activation and ensemble number are similar across tournament days in WT and *Mecp2* KO mice, whereas ensemble similarity diverges between genotypes across days. Furthermore, odor exposure evokes higher mean Ca^2+^ event frequency in WT mice than in *Mecp2* KO mice across all days of the tournament.

### Selective chemogenetic manipulation of vHIP-mPFC activity inversely modulates hierarchy profile in WT and Mecp2 KO mice

We previously showed that selective chemogenetic manipulation of the monosynaptic vHIP-mPFC projection modulates social memory in a pathway-specific and behaviorally selective manner [40], and that hippocampal network hyperactivity in *Mecp2* KO mice due to disinhibition in area CA3 contributes to impaired hippocampal-dependent spatial memory [45, 46, 84, 85]. We hypothesized that hierarchical formation or stability of social ranks in WT mice could be disrupted by chronically activating vHIP inputs to the mPFC using chemogenetics, thereby mimicking the hippocampal hyperactivity observed in *Mecp2* KO mice. Complementarily, reducing hippocampal hyperactivity in *Mecp2* KO mice should alter their hierarchical formation or shift their passive dominance profile, as it did for social memory [40]. To test this, we employed the same intersectional approach as Phillips et al. (2019) [40] to express designer receptors exclusively activated by designer drugs (DREADDs) [86, 87] selectively in mPFC-projecting vHIP neurons, and activated them with the synthetic ligand deschloroclozapine (DCZ) [88–91]. We bilaterally injected adeno-associated virus serotype 8 (AAV8) expressing either the excitatory DREADD hM3Dq into the vHIP of WT mice, or the inhibitory DREADD hM4Di into the vHIP of *Mecp2* KO mice, using a Cre-dependent double-floxed inverse open reading frame (DIO) plasmid [92, 93] at P25–P30 (Fig. 6A). Retrogradely transported CAV2 expressing Cre recombinase (and GFP) was injected bilaterally into the mPFC. Ten days after virus injections, DCZ was provided in the drinking water (1 mg/kg/day) [89, 91, 94]. Control WT and *Mecp2* KO mice were injected with CAV2-Cre in the mPFC and AAV8-DIO-mCherry in the vHIP and administered DCZ.

**Figure 6.**
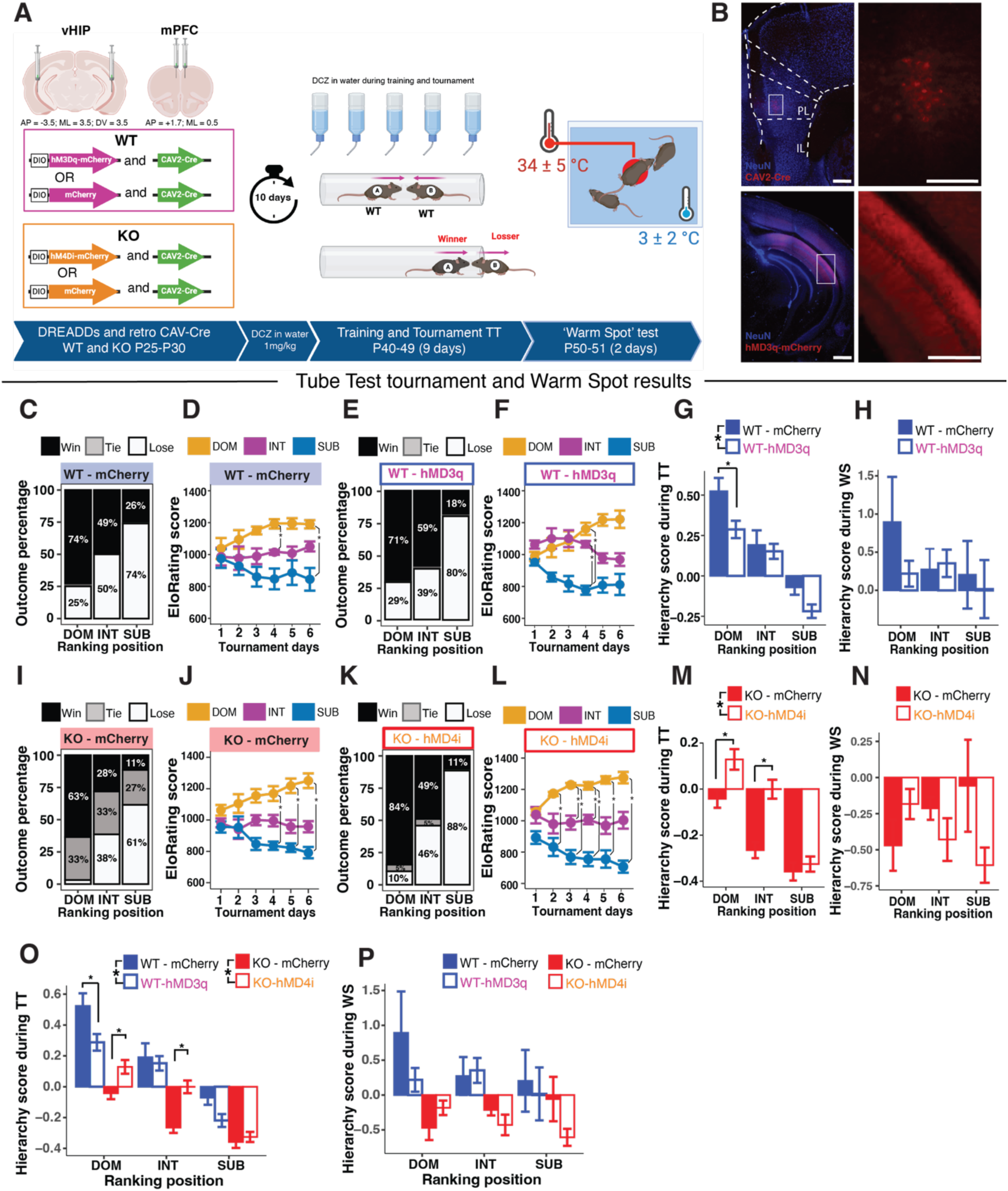
Selective chemogenetic manipulation of vHIP-mPFC activity inversely modulates hierarchy profile in WT and Mecp2 KO mice. (A) Diagram of CAV2-Cre and DREADD bilateral injections followed by posterior tube test tournament and 24 hours after the ‘Warm Spot’. (B) Representative image of injection sites showing CAV2-Cre in mPFC and DREADDs expression in vHIP neurons. Scale bar 200 µm large, 100 µm inset (top), and 500 µm large, 200 µm inset (bottom). (C, I) Cumulative percentages of trial outcomes classifying the 3 male WT-mCherry mice (C) and male *Mecp2* KO-mCherry mice (I) in each cage into dominant (DOM), intermediate (INT), and subordinate (SUB). (D, J) EloRating score in each tournament day for male WT-mCherry mice (D) and male *Mecp2* KO-mCherry mice (J); (E, K) Cumulative percentages of trial outcomes classifying the 3 male WT-hM3Dq mice (E) and male *Mecp2* KO-hM4Di mice (K). (F, L) EloRating score in each tournament day for male WT-hM3Dq mice (F) and male *Mecp2* KO-hM4Di mice (L); (G, H) Hierarchy scores calculated during the tube test (G), and after the Warm Spot test (H) comparing WT-mCherry and WT-hM3Dq showed that WT-mCherry mice exhibited more dominant behaviors after the tournament. (M, N) Hierarchy scores calculated during the tube test (M), and after the Warm Spot test (N) comparing *Mecp2* KO-mCherry and *Mecp2* KO-hM4Di showed that *Mecp2* KO-hM4Di mice exhibited more dominant behaviors after the tournament. (O) Hierarchy scores during the tube test for all four groups. (P) Hierarchy scores during the ‘warm spot’ for all four groups. * *p <* 0.05 with posterior post hoc comparison using Tukeýs HSD.

Both WT (WT-mCherry: n = 12 [4 cages/3 mice]; WT-hM3Dq: n = 12 [4 cages/3 mice]) and *Mecp2* KO (KO-mCherry: n = 12 [4 cages/3 mice]; KO-hM4Di: n = 12 [4 cages/3 mice]) mice were trained for three days to cross the tube in one direction. During the tournament, the percentage of outcomes in both WT and *Mecp2* KO control mice was similar to those in naïve mice (Fig. 1B and 1D). Control WT-mCherry-DOM mice won 73.95%, lost 25%, and tied 1.04%; WT-mCherry-INT mice won 48.9%, lost 50%, and tied 1.04%; and WT-mCherry-SUB mice won 26.04% and lost 73.9% (Fig. 6C). Control *Mecp2* KO-mCherry-DOM mice won 63.54%, lost 3.12%, and tied 33.33%; KO-mCherry-INT mice won 28.12%, lost 38.54%, and tied 33.33%; and KO-mCherry-SUB mice won 11.45%, lost 61.45%, and tied 27% (Fig. 6I). Chronic excitation of the vHIP-mPFC projection in WT mice did not affect tube test outcomes: WT-hM3Dq-DOM mice won 70.8% and lost 29.1%; WT-hM3Dq-INT mice won 59.3%, lost 39.5%, and tied 1.04%; and WT-hM3Dq-SUB mice won 18.75%, lost 80.2%, and tied 1.04% (Fig. 6E). In contrast, chronic inhibition of the vHIP-mPFC projection in *Mecp2* KO mice increased active engagement in competitive interactions, as evidenced by fewer ties: KO-hM4Di-DOM mice won 84.37%, lost 10.41%, and tied 5.2%; KO-hM4Di-INT mice won 48.95%, lost 45.83%, and tied 5.2%; and KO-hM4Di-SUB mice won 11.45% and lost 88.54% (Fig. 6K).

EloRating scores revealed similar hierarchical profiles across groups. A mixed ANOVA showed a significant main effect of rank (F(2, 36) = 82.130, p < .001) and a significant rank × day interaction (F(10, 180) = 17.231, p < .001), but no significant main effect of group (F(3, 36) = 0.102, p = .958) or day (F(5, 180) = 0.140, p = .983), indicating that WT-mCherry, WT-hM3Dq, *Mecp2* KO-mCherry, and *Mecp2* KO-hM4Di mice each established a stable social rank identity throughout the tournament. Notably, *Mecp2* KO-hM4Di mice showed a reduced percentage of ties across all ranks, consistent with increased engagement in competitive confrontations. Chronic excitation of the vHIP-mPFC projection altered the hierarchical score of WT-hM3Dq mice relative to WT-mCherry controls. A mixed ANOVA showed a significant main effect of group (F(1, 570) = 7.551, p < .001; Fig. 6G) and rank (F(2, 570) = 38.565, p < .001), but no group × rank interaction (F(2, 570) = 1.225, p = .294). Post hoc comparisons (Tukey’s HSD) revealed a significant difference between WT-mCherry-DOM and WT-hM3Dq-DOM (diff = −0.236; p = .02), but not between WT-mCherry-INT and WT-hM3Dq-INT (diff = −0.039; p = .997) or WT-mCherry-SUB and WT-hM3Dq-SUB (diff = −0.147; p = .558; Fig. 6G). In contrast, the hierarchical score from the warm spot test showed no significant differences in group (F(1, 18) = 0.740, p = .401), rank (F(2, 18) = 0.543, p = .595), or their interaction (Fig. 6H). These data indicate that WT-hM3Dq mice exhibited lower hierarchy scores in the DOM rank, more subordinate behaviors, and decreased engagement with cagemates, consistent with the predicted effect of mimicking hippocampal hyperactivity in WT mice. The absence of differences in the warm spot test likely reflects the lower number of trials per home cage relative to the tube test tournament.

In contrast, chronic inhibition of the vHIP-mPFC projection in *Mecp2* KO mice altered their grouped hierarchical score relative to KO-mCherry controls. A mixed ANOVA showed a significant main effect of group (F(1, 570) = 24.352, p < .001; Fig. 6M) and rank (F(2, 570) = 49.575, p < .001), and a significant group × rank interaction (F(2, 570) = 4.521, p = .011). Post hoc comparisons (Tukey’s HSD) revealed significant differences between KO-mCherry-DOM and KO-hM4Di-DOM (diff = 0.170; p = .02) and between KO-mCherry-INT and KO-hM4Di-INT (diff = 0.264; p < .01), but not between KO-mCherry-SUB and KO-hM4Di-SUB (diff = 0.032; p = .99). As in WT mice, the warm spot test hierarchical score showed no significant differences in group (F(1, 18) = 1.234, p = .281), rank (F(2, 18) = 0.002, p = .998), or their interaction (Fig. 6N).

Taken together, these observations indicate that chronically reducing vHIP-mPFC activity in *Mecp2* KO mice was sufficient to shift the hierarchy score of DOM mice, increase dominant behaviors, and enhance engagement during social conflicts, mirroring the effects of chemogenetic inhibition on social memory [40]. Chemogenetic manipulation of the vHIP-mPFC projection, increasing excitation in WT mice and decreasing it in *Mecp2* KO mice, thus modulates social hierarchy profiles in opposing directions, underscoring the causal role of this long-range pathway in the social behavioral phenotypes observed in male *Mecp2* KO mice.

## Discussion

In this study, we aimed to determine whether social memory deficits affect social hierarchy formation and social engagement in male *Mecp2* KO mice, and how these behaviors relate to projections from the vHIP to the mPFC. We previously demonstrated that *Mecp2* KO mice exhibit social memory deficits but typical sociability [40]. We ruled out impairments in odor discrimination, indicating that the social memory deficit is not linked to primary sensory deficits in odor perception but rather to higher-order social cognitive processes. *Mecp2* KO mice showed similar engagement in exploring social odors compared to WT mice, indicating preserved discrimination of social stimuli. These findings are consistent with evidence that social recognition and memory enable identification of familiar, novel, or threatening conspecifics [40]. In rodents, social information is primarily conveyed through olfactory cues, with projections from the olfactory bulb to regions including the hypothalamus, hippocampus, and amygdala that process sex, familiarity, and social status [95–98]. Increased neuronal activity in the hypothalamus and medial amygdala, estimated by c-Fos immunostaining, has been reported in male mice exposed to urine odors from conspecifics of different social status [96]. The absence of differences in odor exploration and the intact preference for social versus neutral odors confirm preserved basic olfactory processing of social stimuli in *Mecp2* KO mice.

We then asked whether the previously described social memory deficits [40] affect the formation and maintenance of social rank, considered here as a long-term representation of social identities. Within same-genotype cohorts, both WT and age-matched *Mecp2* KO males formed stable hierarchies, assessed using the tube test and the warm spot test (Figure 1 and Figure 2). However, the quality of interactions differed between genotypes. *Mecp2* KO mice displayed fewer dominant behaviors, reduced motivation for social competition, and less engagement in social conflict. During performance, *Mecp2* KO mice adopted different strategies to resolve conflicts and exhibited reduced hierarchical distance. These features were confirmed in a second conflict assay and were reflected in subordinate behaviors lacking overt dominance traits. This pattern is consistent with a form of passive dominance in which social order is maintained with minimal direct aggression.

Social recognition, memory, and engagement contribute to stable social relationships [99]. Disruption of social memory is associated with altered social behavior in psychiatric conditions [12, 100, 101]. Social memory has also been linked to activity in reward-related [102] and memory-related brain circuits [103], and animals will work to access social interaction as a motivational stimulus [104]. The reduced aggression and social engagement observed in *Mecp2* KO mice (Figure 1 and Figure 2) are therefore consistent with decreased social motivation rather than impaired social recognition per se. Altered mesolimbic function, implicated in social motivation, may contribute to this phenotype [105].

Prior studies from our and other laboratories have identified an altered excitation/inhibition balance in *Mecp2* -deficient mice in regions relevant to social memory, including hyperactivity in areas CA3 and CA1 of the ventral hippocampus [45, 46, 85, 106] and lower activity and hypoconnectivity of excitatory neurons in the prelimbic mPFC [40, 107–109]. This imbalance suggests that the monosynaptic vHIP–mPFC projection may contribute to social rank expression, as it does for social memory [40]. Chronic chemogenetic stimulation of this pathway in WT mice with the excitatory DREADD hM3Dq shifted dominance hierarchy (Figure 6), whereas chemogenetic silencing in *Mecp2* KO mice with the inhibitory DREADD hM4Di increased dominant behaviors. These findings are consistent with evidence that vHIP–mPFC signaling regulates anxiety-related behaviors and aversive representations [110], and that cortical projections modulate social approach and avoidance [111]. Together with previous work demonstrating rescue of social memory deficits in *Mecp2* KO mice after vHIP–mPFC inhibition [40], these data support a role for this projection in the modulation of social behavior.

Although altered excitation/inhibition balance in the mPFC has been linked to sociability deficits in some mouse models of ASD [15, 78, 112], other regions also contribute to social behavior. The CA2 hippocampal area has been implicated in social memory and aggression [43, 113–116]. In mouse models for Rett syndrome based on *Mecp2* deletion, altered plasticity of perineuronal nets in CA2 has been reported [103, 117]. Dorsal CA2 contributes to social novelty recognition and aggression [42, 43, 113, 118]. Activity of dorsal CA2 neurons is modulated during both non-aggressive and aggressive interactions, and *Avpr1b* signaling from PVN projections promotes aggression [113, 118]. In *Mecp2* KO mice, resistance to synaptic plasticity in dorsal CA2 due to perineuronal net alterations has been described, and matrix degradation restores plasticity [117, 119]. Additional alterations in models for Rett syndrome include changes in cortico–lateral amygdala synapse maturation [120], reduced amygdala volume [121], altered stress-related gene expression in the PVN, amygdala, and BNST [122], and gene expression changes in the hypothalamus and cerebellum [123]. These regions contribute to social interaction, social memory [76, 120, 124, 125], and social rank stability.

Social rank formation involves multiple brain regions, including the striatum [126], basolateral amygdala [127], lateral hypothalamus [59], anterior cingulate cortex [128, 129], and mPFC [53, 54]. Neuronal population activity in the mPFC predicts social rank [59]. Given the previously reported mPFC hypoactivity in *Mecp2* KO mice [40], altered cortical processing likely contributes to the reduced hierarchical differentiation observed here.

Calcium imaging from excitatory mPFC neurons in WT mice identified neurons with either increased or decreased activity during social interaction relative to baseline (Suppl. Fig3). Socially sensitive neurons were present in both WT and *Mecp2* KO mice during the warm spot test, with similar proportions of positive, negative, and non-responsive cells (Figure 3). However, mPFC pyramidal neurons in *Mecp2* KO mice exhibited lower activity, reflected in their smaller amplitude and occurrence, during both baseline and competition. No differences in activity were observed as a function of an established rank.

Theoretical frameworks propose that the mPFC integrates social information with valence and salience signals [59, 130]. Synchronous oscillations between the vHIP and mPFC support working memory [131] and are driven by monosynaptic glutamatergic projections from ventral CA1/subiculum to mPFC pyramidal neurons and interneurons [74, 132–134]. We therefore examined ensemble coactivation during social events. Neuronal ensembles, coactive groups proposed to encode memory and behavior [69–71], showed reduced functional coactivation in *Mecp2* KO mice during social competition (Figure 4). In contrast, ensemble coactivation during exposure to urine odors did not differ between genotypes, consistent with intact odor discrimination. Given the convergence of inputs to the mPFC [74, 75] and its integrative role in emotion and cognition [15, 40, 60, 76, 77], reduced ensemble coordination may reflect impaired integration of social information rather than a sensory processing deficit.

Reports of sociability in mouse models for Rett syndrome vary depending on the specific genotype, testing conditions, and genetic background [135–139]. In the *Mecp2* KO mouse line used here, which carries a deletion of exon 3 (Jaenisch line), sociability is comparable to WT mice, but social memory is impaired [40]. Here, we assessed social dominance as a long-term memory of social identity. Prior tube test studies in *Mecp2* mouse models report mixed findings [136, 139], potentially due to differences in *Mecp2* mutations, genetic background, and testing or housing conditions. Across cohorts, WT mice displayed more dominant behaviors than *Mecp2* KO mice, independent of age. Lower aggression in *Mecp2* models has been reported previously [137, 140]. Altered serotonin, oxytocin, or vasopressin signaling may contribute to this phenotype [99, 141–143]. Conversely, overexpression of human *MECP2* in mice modulates aggression in a background-dependent manner [144]. Other mouse models for neurodevelopmental disorders also show lower dominance, including *Ext1* conditional KO mice [145], mice exposed to environmental toxins [146], *Fmr1* KO mice [147], and *Dvl1*-deficient mice [148]. Hyperserotonemia in *SERT* Ala56 mice increases submissive behaviors [149]. In contrast, increased dominance has been reported in *Dlgap2* KO mice [150] and in some studies using *Fmr1* KO mice [151, 152], indicating model-dependent effects on social hierarchy.

We assessed hierarchy formation within same-genotype groups, because social rank is an emergent property of repeated social interactions [130]. Mixed genotype or unfamiliar pairings may instead reflect individual dominance traits rather than socially learned hierarchy. Group housing supports long-term social memory after brief interactions [153], suggesting that social context influences dominance outcomes.

In summary, male *Mecp2* KO mice do form stable hierarchies despite their social memory deficits, but display reduced social engagement and dominance behaviors during competition-based tasks. The vHIP–mPFC projection modulates this social competition, and altered circuit hyperactivity in this pathway contributes to the observed behavioral differences. These findings support a circuit-level contribution of vHIP–mPFC signaling to the social behavior alterations observed in *Mecp2* deficiency.

## Materials and Methods

### Animals

Female mice from the Jaenisch strain with a deletion of exon 3 of the *Mecp2* gene (Mecp2tm1.1Jae) [154])were obtained from the *Mutant Mouse Resource & Research Center*, and a colony established at the University of Alabama at Birmingham (UAB) by crossing them with male WT C57/BL6 mice. All mice used in these experiments were male hemizygous Mecp2tm1.1Jae (*Mecp2* KO), and controls were age-matched male WT littermates. All experimental subjects were 5–6 weeks of age (P35–P42) at the time of surgery, behavioral testing, and *in vivo* Ca^2+^ imaging. At this age, male *Mecp2* KO mice do not exhibit overt Rett-like symptoms, such as hypoactivity, hind limb clasping, resting tremor, or reflex impairments, which typically appear between P45 and P60 [40, 155]. Mice were genotyped and weaned at 3 weeks of age and maintained on a 12:12 h light/dark cycle (lights on 6:00–18:00 h) with food and water *ad libitum*. All procedures and housing followed recommendations of the *Committee on Laboratory Animal Resources* of the National Institutes of Health (NIH). Experimental protocols were approved by the *Institutional Animal Care and Use Committee* (IACUC) of UAB.

### Behavioral tests

#### Odor habituation-dishabituation test

During the light phase (between 9:00 AM and 4:00 PM), male WT (n = 16) and *Mecp2* KO (n = 16) mice aged P30–P40 performed the odor discrimination test. Two days before testing, a wooden cube (2 cm³) was placed inside the home cage and remained there until the test day. Pieces of chocolate cookies were also placed in the home cage two days before testing. On the test day, each mouse was individually subjected to a habituation–dishabituation procedure in the center of an empty cage (40 cm × 30 cm) without bedding. This procedure was adapted from [156]. Mice were first allowed to freely explore the cage for 10 min with an unscented wooden cube, identical to the one placed in the home cage, positioned at least 10 cm from any wall. After this familiarization period, the cube was removed for 1 min (inter-trial interval, ITI). A new wooden cube scented with distilled water was then placed in the same location for 2 min and removed for a 1-min ITI. This procedure was repeated three times (3×), constituting the habituation phase. Following the final ITI, a new wooden cube scented with the test odor (either social [unknown male urine] or non-social [chocolate odor], 1:400 dilution) was presented once (1×), corresponding to the dishabituation phase. Each mouse was exposed to both social and non-social stimuli in a balanced order: half of the WT and *Mecp2* KO mice were first exposed to the social stimulus, and the remaining half were first exposed to the non-social stimulus. After completing four trials (3× habituation and 1× dishabituation for the first odor), mice rested for 5 min in the experimental cage without a cube. A second habituation phase with distilled water (3×) was then conducted, followed by a dishabituation trial (1×) with the second odor. Odor exploration time and tracking were automatically quantified using EthoVision XT software (version 16, Noldus) from offline video recordings. Exploration was defined as the mouse’s nose being within approximately 1.5 cm of the cube edges (odor zone).

#### Social hierarchy from the tube test

Hierarchies were established among mice housed in groups of three of the same genotype. Groups were randomly formed and matched for age. Two independent cohorts were trained and tested using the standard tube test [52, 54]. In the first cohort (N = 30; WT = 12 and *Mecp2* KO = 18; P40), training followed the protocol described by Fan et al. (2019) [52], consisting of three days of individual tube crossing, followed by a round-robin tournament conducted over six days. To rule out potential training deficits related to altered pain sensitivity—given that *MECP2* mutations in Rett individuals have been associated with altered pain sensitivity [157], and animal models have reported changes in mechanical sensitivity and pain [158, 159]—a second cohort was tested. The second cohort (N = 36; WT = 18 and *Mecp2* KO = 18; P25–P30) was trained to cross the tube without the constant tail handling during each training trial and without the push-encouraging used with the first cohort. The six-day round-robin tournament phase of the tube test was identical between cohorts. Following the tournament, both cohorts were subjected to the warm-spot test. In the first trial, mice were individually exposed to an open field with a cold floor containing a single warm spot. Twenty-four hours later, mice were exposed as a group to the same open field with a cold floor and one warm spot, as originally described by Zhou et al. (2017) [54].

##### Training

The first cohort of mice was trained individually for three days. A Plexiglas tube (30 cm length, 2.8 cm internal diameter) was positioned at the center of a large arena (75 cm × 30 cm). During the first 2 min, each mouse was allowed to freely explore both the tube and the arena. After this habituation period, the mouse was gently lifted by the tail and placed at one end of the tube, and released once it entered the tube. If the mouse stopped or attempted to retreat while crossing, it was gently touched on the tail or back with a plastic stick. A trial was considered complete when the mouse’s paws fully exited the tube. The procedure was then repeated from the opposite end. Each mouse crossed the tube ten times per day (five trials from each side). Trial duration was automatically quantified using EthoVision XT software (version 16, Noldus), and the number of pushes was manually scored from offline video recordings. Previous studies in *Mecp2* KO mouse models have reported altered pain sensitivity and hypersensitivity [158, 160, 161]. In the first cohort, *Mecp2* KO mice required a higher number of encouragements during training. To control for potential effects of repeated handling and touching on subsequent social hierarchy assessment, a second cohort of age-matched WT and *Mecp2* KO mice was trained with minimal manipulation for four days. In this second cohort, mice were gently placed into a small Plexiglas cage (8 cm length, 9 cm width, 9 cm height) connected to the Plexiglas tube (30 cm length, 2.8 cm diameter). During the first 2 min, each mouse explored the tube and the attached small cage. A metallic door then blocked the tube entrance in the small cage where the mouse was located. Ten seconds later, the door was removed, allowing the mouse to enter the tube. Mice were given up to 3 min to cross the tube. Once the mouse entered the tube, a metallic door prevented retreat. After the mouse entered the opposite small cage, another metallic door blocked the tube entrance for 10 s. As in the first cohort, mice performed ten trials per day (five from each side). If 3 min elapsed without entry into the tube, the lid of the small cage was opened, and the experimenter’s hand was slowly introduced to guide the mouse into the tube. Trial duration was quantified using EthoVision XT software (version 16, Noldus). As there were no differences in crossing time or performance during the tube test tournament between cohorts, behavioral data related to social hierarchy were pooled for analysis.

##### Round-robin tournament

after training, mice competed pairwise in a round-robin tournament using the standard tube test [52] over six consecutive days to establish hierarchy within each home cage. Each daily tournament consisted of two round-robin sequences: in the first, each pairwise combination of animals competed once; in the second, the same pairings were repeated in reversed order. Thus, each day included six competitions per cage (e.g., A–B, A–C, B–C, followed by B–A, C–A, C–B). During each competition, two mice were gently lifted by the tail, placed at opposite ends of the tube, and released simultaneously once both entered the tube. Mice were allowed up to 3 min to compete to exit through the opposite end, typically resulting in a winner and a loser [52, 55, 56]. A mouse was classified as a winner if it pushed the opponent out of the tube or remained inside while the opponent retreated. The mouse exiting the tube was classified as the loser. In some trials, mice passed each other without retreating or pushing (negotiated crossing), or both remained inside the tube for the full 3 min without resolution; these outcomes were classified as ties. Absolute social rank was determined from the overall percentage of wins, losses, and ties across the six tournament days (Fig. 2A), classifying mice within each cage of three as dominant (DOM), intermediate (INT), or subordinate (SUB). In addition to daily outcomes (wins, losses, ties), latency to resolve the contest and body interactions were quantified from offline video recordings using EthoVision XT software (version 16, Noldus). Frame-by-frame behavioral analysis inside the tube was performed using BORIS software [52, 162]. The following behaviors were manually annotated: (a) *pushing*, defined as pressing and shoving the head under another mouse to gain territory; (b) *pushing back*, defined as a counter-push following an initial push; (c) *resistance*, defined as maintaining position while being pushed; (d) *retreat*, defined as backing out voluntarily or in response to a push; (e) *chasing*, defined as pursuing a retreating mouse; and (f) *stillness*, defined as remaining immobile during the confrontation. *Pushing*, *pushing back*, *resistance*, and *chasing* were categorized as aggressive behaviors, whereas *retreat* and *stillness* were categorized as submissive behaviors.

Hierarchical distance between mice was quantified using the EloRating score [129, 163], a probabilistic dominance model that continuously updates individual ratings based on wins and losses. Expected outcomes result in smaller rating changes than unexpected outcomes. The Elo rating complemented ordinal rank (based on percentages of wins, losses, and ties) by quantifying the magnitude of dominance differences between individuals and distinguishing strong from weak hierarchical relationships, and it allows parametric statistical analysis across multiple response variables [129]. Elo ratings were calculated using the EloRating R package [163]. To further quantify hierarchical position, we calculated a composite hierarchy score based on dominant behaviors, similar to previous reports [67, 164]. Z-scores were calculated for the total number of wins during the pairwise tournament and for *pushing*, *pushing back*, *resistance*, and *chasing* behaviors using the formula:

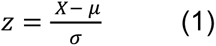

where X represents the individual value, μ the mean, and σ the variance. The hierarchy score was then calculated as:

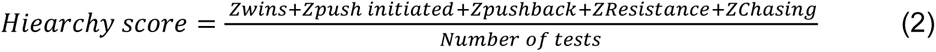

Higher hierarchy scores indicate a greater expression of dominant behaviors across the tournament, whereas scores near zero or negative values indicate subordinate behavior or reduced dominance.

#### Social hierarchy from the warm spot test

Complementary to the tube test, and to identify additional forms of social interactions, a warm spot (WS) test was performed 24 h after the final day of the tube test tournament. The WS test was adapted from Zhou et al. (2017) [54] and consisted of two consecutive days. On the first day, mice were individually placed for 10 min in an open arena (25 cm × 30 cm) that had been pre-cooled on ice, resulting in a floor temperature of approximately 1–3 °C. Beneath the floor, an adhesive polyimide heater plate connected to a digital temperature controller maintained the temperature of the warm spot at approximately 37–40 °C. The warm spot had a diameter of 4 cm, allowing occupancy by only one mouse at a time. During this session, mice explored the cold arena and, in most cases, located the warm spot, where they remained for extended periods. In a small number of cases in which mice had not located the warm spot after 5 min, the experimenter gently placed the mouse onto the warm spot, where it remained thereafter. On the second day, all mice from the same home cage were reintroduced together into the cold arena with the warm spot positioned in the same location, and behavior was recorded for 10 min. The time spent on the warm spot and the frequency of warm spot access were quantified. Both days were recorded using EthoVision XT software (version 16, Noldus). For the group session on the second day, behavior was manually scored using BORIS software by an observer different from the experimenter. This approach was necessary because automatic tracking resulted in errors that resulted in swapping of identities between mice, preventing reliable individual identification. To facilitate offline identification, mice were marked on the back with a small white paper tag. In addition to time and frequency on the warm spot, the following behaviors were manually annotated: *grooming* (frequency and duration), *stillness* or *immobility*, *pushing*, *jumping*, social exploration (*nose-to-nose*, *nose-to-anogenital*, *nose-to-back*), *chasing* or *following*, and *climbing* over a cagemate.

#### 4-Chamber social arena for social preference and social memory

The experimental apparatus consisted of a hexagonal central chamber connected to three peripheral chambers by transparent, perforated partitions [41]. Test mice were allowed to move freely within the central chamber, while different stimuli were placed in the three side chambers: a non-social object (toy mouse), a cage mate matched for sex and age, and a novel mouse also matched for sex and age. The perforated partitions allowed the exchange of whisker and olfactory cues between the test mouse in the central chamber and the stimuli in the side chambers. The experimental design included a two-day acclimation period during which mice were familiarized with the central chamber of the four-chamber social arena for 5 min per day. Following habituation, two sequential trials were conducted. In the first trial (habituation phase), mice were allowed to explore the central chamber for 5 min. In the second trial (sociability phase), a toy mouse and a cage mate were placed in separate side chambers, and the test mouse was allowed to explore for 5 min to assess preference for the live stimulus (sociability). All trials were conducted under low red light and recorded from above at 30 frames per second using an infrared-sensitive camera (scA780-54gm, Basler) controlled by EthoVision XT17 (Noldus). Time spent within 2.5 cm of each partitioned area was quantified using EthoVision.

### *In vivo* Ca^2+^ imaging

#### Surgeries to express Ca^2+^ sensors and implant GRIN lenses

The first cohort of male mice (P45) were anesthetized with 3% isoflurane vapor in 100% oxygen and then placed in a stereotaxic frame (Kopf Instruments, Tujunga, CA) maintained under 2% isoflurane during surgery. Body temperature was monitored with a rectal probe (Physitemp, NJ) and maintained using a temperature-controlled heating pad (TCAT-2AC, Physitemp, NJ). A midline scalp incision was made to expose the skull, and a small craniotomy was performed over the right hemisphere using a dental drill. Injections were performed using a 2.5 µL Hamilton syringe (Hamilton Company, Reno, NV) mounted on a microsyringe pump (UMP3 UltraMicroPump, Micro4; World Precision Instruments, Sarasota, FL) at a rate of 0.30 µL/min. An adeno-associated virus, serotype 9 encoding GCaMP6s under the *CamkIIα* promoter (AAV.CamKII.GCaMP6s.WPRE.SV40; titer 1.0 × 10¹³ vg/mL; Addgene, Watertown, MA) was injected (300 nL) into the prelimbic prefrontal cortex at the following coordinates: AP +1.7 mm, ML +0.45 mm, DV −2.0 mm. The needle remained in place for 5 min after completion of the injection before being slowly withdrawn. The incision was then closed with surgical glue. Mice were allowed to recover for at least 3 weeks. For GRIN lens implantation, mice underwent a second surgery under the same anesthetic conditions (3% isoflurane induction, maintained at 2%). A craniotomy (∼2 mm diameter) was performed over the previous injection site. Using a blunt needle connected to vacuum aspiration and continuous sterile saline perfusion, approximately 1.5 mm of overlying cortical tissue was gently aspirated. After hemostasis was achieved, a GRIN lens attached to a base plate (ProView™ Integrated Lens [PIL], 4 mm length, 1 mm diameter; Inscopix, Palo Alto, CA) was slowly lowered stereotaxically into the prelimbic cortex mPFC to a depth of −2.0 mm at a 10° angle. The base plate was secured to the skull using dental cement. After the dental cement hardened, ketoprofen (10 mg/kg; AlliVet, St. Hialeah, FL) was administered as an analgesic. Animals were allowed to recover for at least 2 weeks before assessing GCaMP expression and proceeding with behavioral testing in the 4-chamber social arena.

To avoid multiple surgeries prior to miniscope imaging, we followed the procedures described by Jackman et al. 2018 [61]. Briefly, silk fibroin (40% of the mixture; 1.0 µL; Sigma Aldrich) was combined with AAVs (60% of the mixture; 1.5 µL of AAV.CamKII.GCaMP8m.WPRE.SV40; titer 1.0 × 10¹³ vg/mL; Addgene, Watertown, MA). A total of 1.5 µL of the mixture was applied to the surface of a GRIN lens (1.0 mm diameter) in a single application from above and allowed to air dry. Following implantation, the silk matrix gradually released AAVs, resulting in localized transduction of cells surrounding the GRIN lens. During surgery, approximately 1.5 mm of cortical tissue was aspirated, and the pre-coated GRIN lens was lowered stereotaxically to the final target depth (DV −2.0 mm) at a 10° angle. Two weeks after surgery, mice were evaluated for GCaMP8m expression. Pilot experiments indicated that pre-coating GRIN lenses with the AAV–silk fibroin mixture did not affect GCaMP8 fluorescence intensity or the amplitude of ΔF/F transients reflecting intracellular Ca²⁺ signals in mPFC pyramidal neurons (Suppl. Fig. 4). Following these pilot studies, both WT and *Mecp2* KO mice were implanted at P25–P30. Ten days after surgery, mice began training in the tube test. For head-fixed imaging, the procedure was similar to the pre-coated GRIN lens surgeries. After implantation of the pre-coated GRIN lens, a custom head-fixation device was positioned over the posterior skull and secured with dental cement. Two weeks after surgery, mice were trained to walk in a wheel apparatus for head-fixed imaging.

#### Miniscope imaging

Mice were habituated to the weight of the miniscope using a dummy miniscope (2 g; Inscopix, Palo Alto, CA) for at least 2 days before imaging. Following the tube test tournament, on the first day of the warm spot test (habituation session), mice were fitted with the miniscope and placed in the arena. Imaging was performed using a head-mounted miniaturized microscope (i.e. miniscope; nVoke™, Inscopix, Palo Alto, CA) equipped with an integrated 475-nm LED. Images were acquired at 20 frames per second with the excitation LED power set to 0.2–0.3 mW while mice moved freely in the arena. Gain and focus parameters were determined individually for each mouse and maintained constant across day zero and day one of the warm spot test. The image data acquisition system (IDAS, Inscopix, Palo Alto, CA) was synchronized with EthoVision XT17 software via a transistor–transistor logic (TTL) pulse (5 mV) to simultaneously initiate imaging and behavioral movies. Data were acquired continuously for the full 10 min duration of the warm spot test.

#### Data analysis

Calcium imaging data were analyzed as previously described. During offline processing, recordings were spatially downsampled by a factor of two, bandpass filtered, and motion corrected using default parameters using *Inscopix Data Processing Software* (version 1.9.5; IDPS, Inscopix, CA). Individual traces of jGCaMP8m fluorescence were extracted using *Constrained Nonnegative Matrix Factorization* for microendoscopic data (CNMFe) [165]. The following parameters were used for CNMFe extraction: minimum correlation = 0.8; minimum peak-to-noise ratio (PNR) = 320; Gaussian filter = 3; gSiz = 20; gSig = 10. Extracted components were manually inspected, and non-neuronal signals were excluded. Individual event amplitude and frequency were quantified using custom-written MATLAB scripts.

##### jGCaMP8m signal processing

To identify neurons responsive to specific behaviors, we followed a procedure similar to that described by Patel et al. 2022 [64]. After CNMFe extraction, individual traces of jGCaMP8m ΔF/F signals were binned into 1 s intervals. Signals were Z-normalized using the 3 s preceding behavioral onset as baseline. Traces exceeding ± 2 standard deviations (SD) from the Z-normalized baseline were classified as positively (above + 2 SD) or negatively (below − 2 SD) responsive. As a control, the average trace was compared to the same trace after random permutation. In Fig. 3O–V, green bars beneath the traces indicate time points reaching statistical significance, with p-values calculated using false discovery rate (FDR) correction. FDR was computed from point-by-point t-tests and adjusted for multiple comparisons using the criterion P(k) < 0.05 × k/2000 [67].

##### Synchronization of behavioral annotations with jGCaMP8m imaging

Behavioral videos acquired during the warm spot test were manually annotated and synchronized with jGCaMP8m recordings using *Behavior Ensemble and Neural Trajectory Observatory* (BENTO) software [63] (http://github.com/neuroethology/bentoMAT). BENTO is a MATLAB-based graphical user interface that enables frame-by-frame behavioral annotation, similar to procedures implemented in BORIS software. In addition, BENTO allows simultaneous visualization of neural calcium recordings, multiple video streams, and manually annotated behavioral events. Exploratory analyses of neural–behavioral synchronization indicated that subsets of neurons increased activity at the onset of specific behaviors, whereas another subset exhibited reduced signal amplitude at behavioral onset. Initial exploratory correlations between calcium activity and behavior were performed within BENTO. The platform also provides tools for basic neuronal data analyses, including k-means clustering and dimensionality reduction.

##### Detection of neuronal ensembles based on correlated activity

To identify neuronal ensemble activity, we followed the procedure described by Perez-Ortega et al. (2021, 2024) using custom MATLAB code (Xsembles2P) developed for initial analysis of calcium imaging data (https://github.com/PerezOrtegaJ/Xsembles2P). Prior to this analysis, putative spikes (events) were inferred from CNMFe-extracted calcium traces using deconvolution with a signal-to-noise ratio (SNR) threshold of 3 and the OASIS algorithm (Online Active Set method to Infer Spikes) [57], generating a binarized signal (Fig. 4A). The binarized activity was organized into a raster matrix to extract the functional neuronal network. A filtering step was applied to remove spikes from neurons lacking functional connections within each column vector. After preprocessing, hierarchical clustering was performed on all column vectors using the *Jaccard* similarity coefficient combined with *Ward* linkage (Fig. 4A). The optimal number of clusters was determined by identifying the highest local contrast index. Clusters were classified as ensembles when the similarity between vectors reached statistical significance (p < 0.05). Significance was determined by comparing the average within-cluster similarity to the distribution of average similarity values obtained from randomly generated clusters across 1000 iterations. For each significant ensemble, timestamps of ensemble activity were extracted (Fig. 4E–H).

#### Chemogenetic manipulation with DREADDs and DCZ

We followed the intersectional conditional approach to express DREADDs in mPFC-projecting ventral CA1 neurons described in Phillips et al. 2019 [40]. Both WT and *Mecp2* KO mice received bilateral injections of retrogradely transported CAV-2-Cre (Biocampus, Institute of Molecular Genetics, Montpellier, France) into the mPFC (300 nL per site; AP +1.7 mm, ML +0.5 mm, DV −2.0 mm). *Mecp2* KO mice were bilaterally injected in the vHIP (AP −3.5 mm, ML +3.5 mm, DV −3.5 mm) with 500 nL per site of AAV8-hSyn-DIO-mCherry or AAV8-hSyn-DIO-hM4Di(Gi)-mCherry to express an inhibitory DREADD. WT mice received bilateral injections of AAV8-hSyn-DIO-hM3Dq(Gq)-mCherry (500 nL per site) to express an excitatory DREADD. Deschloroclozapine (DCZ; Sigma Aldrich) was first dissolved in saline (5 mg in 5 mL) and subsequently diluted into 200 mL of water containing 5 mM saccharine (Sigma Aldrich) [88–91]. Mice had *ad libitum* access to this solution starting at postnatal day 34 (P34) via a standard drinking bottle. DCZ treatment was maintained until euthanasia, which occurred 3–5 days after completion of the warm spot test.

#### Histology and immunohistochemistry

To confirm GCaMP expression and GRIN lens placement, mice were anesthetized with ketamine (100 mg/kg) and xylazine (10 mg/kg) and transcardially perfused with 1× phosphate-buffered saline (PBS), followed by ice-cold 4% paraformaldehyde (PFA) in 1× PBS. Brains were extracted and postfixed overnight in 4% PFA. Coronal sections (60 μm) were cut using a vibratome (PELCO 100, model 3000; Ted Pella Inc., Redding, CA). For direct mounting, sections were placed on glass slides and coverslipped with Vectashield mounting medium (Vector Biolabs, Malvern, PA). For immunohistochemistry, free-floating sections were stored at 4°C in 1× PBS containing 0.01% sodium azide. Sections were permeabilized in 0.25% Triton X-100 for 15 min and blocked for 1 h in a solution containing 0.01% sodium azide, 2% bovine serum albumin (BSA), 0.1% Triton X-100, 2 M glycine, and 10% goat serum in 1× PBS. Primary antibodies were diluted in antibody diluent (0.01% sodium azide, 2% BSA, 0.1% Triton X-100, and 5% goat serum in 1× PBS) and incubated for 36 h at room temperature. After three washes in 1× PBS (5 min each), sections were incubated with secondary antibodies diluted in the same diluent for 4 h at room temperature. Sections were then washed three times (5 min each) in 1× PBS and mounted with Vectashield mounting medium (Vector Biolabs, Malvern, PA). For primary antibodies requiring signal amplification (e.g., mCherry or GFP), an additional biotin amplification step was included after primary antibody incubation. Sections were incubated with biotinylated anti-host secondary antibodies (Vector Biolabs, Malvern, PA; 1:500 dilution) in antibody diluent for 2 h, washed three times in 1× PBS (5 min each), and then incubated with streptavidin-conjugated fluorophores (Alexa-405, Alexa-488, Alexa-594; Thermo Fisher Scientific, Waltham, MA; 1:500 dilution) for 4 h. After three additional washes (5 min each), sections were mounted with Vectashield mounting medium.

#### Statistical analyses

For parametric data, comparisons between two groups were performed using Student’s t-tests. For comparisons involving multiple groups or factors, one- or two-way analysis of variance (ANOVA) was applied, with group or rank as factors. Repeated-measures ANOVA was used for behavioral experiments when appropriate. Significant main effects or interactions were followed by post hoc pairwise comparisons using Bonferroni correction or Tukey’s HSD test. For non-parametric data, Mann–Whitney or Kolmogorov–Smirnov tests were used as appropriate. Statistical significance was set at p < 0.05. Sample sizes were determined a priori using power analysis in G*Power software [166] to achieve at least 80% power. The specific statistical tests and sample sizes for each experiment are provided in the main text and corresponding figure legends.

## AUTHOR CONTRIBUTIONS

CA-T designed and performed experiments, analyzed data, and wrote the manuscript; JJT and JJD provided key reagents and their optimization; JP-O provided custom written analysis code and perform some data analysis; AS performed some data analysis; SC performed some imaging sessions and data analysis; LP-M designed experiments, analyzed data, and wrote the manuscript.

## ACKNOWLEDGEMENTS

We are grateful to Dr. Destynie Medeiros, Mr. Chang Li, and Drs. Wei Li and Xin Xu for training and discussions throughout these studies, and to Mr. Yijian Zhang for mouse colony maintenance. We thank Drs. Alejandro Osorio-Forero (Netherlands Institute for Neuroscience) for valuable insights and help writing some parts of the MATLAB code for calcium imaging analysis, and Luis Rosas-Vidal (Northwestern University) for sharing MATLAB code for calcium imaging analysis. Funding was provided by a Fulbright Scholarship (CAT) and NIH grant MH-118563 (LPM).

---

**Supplemental Figure 1. Male *Mecp2* KO mice have typical odor discrimination**

Because olfactory discrimination is necessary for social identification in mice, we tested social and non-social odor discrimination in male *Mecp2* KO mice and compared it to that of male WT mice with the “odor habituation/dishabituation test” using wooden cubes impregnated with either water, a non-social stimulus (chocolate) and of a social stimulus (urine). (**A**) Design of the “odor habituation/dishabituation test”. **(B)** Representative tracks of a male WT mouse (top) and a male *Mecp2* KO mouse (bottom) during exploration of a wooden cube (red indicates more time spent at that location). **(C Top)** Frequency of close interactions with wooden cube and exploration time of wooden cube **(C Bottom)** during the 3 “habitation” trial with a neutral cube (water), and the final “dishabituation” trial with either a social or non-social cube (shaded area). (**D**) Similar distance traveled during odor discrimination. (**E**) Similar velocity during odor discrimination. * *p <* 0.05 adjusted after Bonferroni correction.

**Supplemental Figure 2. Additional information on the warm spot test**

**(A)** Design and measurements of the ‘warm spot’ arena. Heatmap shows a representative location of a WT mouse during 10 mins. **(B)** Differences in the distance traveled during day of training in the ‘warm spot’ comparing WT and Mecp2 KO mice. WT mice showed more gross activity than *Mecp2* KO. (C) Differences in the speed during day of training in the ‘warm spot’ comparing WT and Mecp2 KO mice. WT mice moved more faster than *Mecp2* KO. (D) Example of an ethogram during ‘warm spot’ test showing behaviors scored and duration. (E) Diagram of behaviors scored during the ‘warm spot’ test. Top left. Frequency on ‘warm spot’, and pushing. Left second row, chasing. Left third row: jumping and resting/cuddling, Left forth row, grooming. Right column, showing representation of different social interactions, nose-to-nose, nose-to-body, nose-to-rear, and over cagemate.

**Supplemental Figure 3. Socially sensitive neurons in mPFC that were active during social interactions were identified by in vivo Ca^2+^ imaging with head-mounted miniscopes**

(A) Diagrams of head-mounted miniscope with GRIN lens in the mPFC. (B) Example of GRIN lens placement aimed at jGCaMP6s-expressing excitatory neurons (CaMKII promoter). (C) Unprocessed image of jGCaMP6s fluorescence. (D) The same field-of-view (FOV) as in C after motion correction and ICA-PCA processing, showing neurons from where jGCaMP6s signals were obtained. (E) 4-chamber arena: test mouse in the center chamber has the choice to approach 3 side chambers separated by perforated partitions. Left: habituation. Right: novel mouse in top left side chamber. (F) Example of traces of jGCaMP6s signals (% dF/F) during habituation (left of vertical dotted line) and social exploration (right of vertical dotted line). Social sensitive inhibited (as known as Social-OFF, [60]) neurons (blue traces) show fewer Ca2+ transients when the mouse approaches the chamber with a novel mouse (gray shading) than at other times. Social sensitive activated (as known as Social-ON [60]) neurons (red traces) show more Ca2+ transients when the mouse approaches the chamber with a novel mouse (gray shading) than at other times.

**Supplemental Figure 4. Social responsive neurons in mPFC were similar between *Mecp2* KO and WT mice, and similar dynamics between dominant and submissive social ranking**

(A). Pre-coating GRIN lenses with a mixture of AAVs and silk fibroin did not affect neither GCaMP6 fluorescence intensity nor the amplitude of its dF/F transients reporting intracellular Ca2+ levels in mPFC pyramidal neurons. Diagram of coating a GRIN lens with a mixture of AAV-JGCaMP6s and silk fibroin (Jackman et al., 2018), and the comparison with a GRIN lens pre-coated with silk only to image jGCaMP6s-expressing neurons after standard AAV injections. (B) Representative FOVs from the mPFC of WT mice using a GRIN lens pre-coated with silk and jGCaMP6s (top), and silk only (bottom), after 2 weeks (left) and 6 weeks (right) from lens implantation; all images are maximum time projections of dF/F, after motion correction and CNMFe processing. (C and D) Representative jGCaMP6s traces (% dF/F) during habituation and social interaction test. (C) traces are from a mouse with jGCaMP6s/silk GRIN, and, (D) traces from a mouse with silk-only GRIN and AAV-jGCaMP6s injection. (E) Average dF/F z-scores during different social behaviors showing average of traces classified as positive responsive in WT and *Mecp2* KO mice, FDR (represented as a green line) showed statistically significant differences in those points during nose-to-body interactions. (F) Average dF/F z-scores classified as negatively responsive in WT mice and in *Mecp2* KO, FDR (represented as a green line inside the plot) showed statistically significant differences in those points during over-cagemate interactions. (G) Average dF/F z-scores from positive and negative responsive neurons in WT-DOM and *Mecp2*-DOM across different social behaviors. (H) Average dF/F z-scores from positive and negative responsive neurons in WT-SUB and *Mecp2*-SUB.

**Supplemental Figure 5. Pyramidal mPFC neurons showed higher correlation scores in WT mice than in *Mecp2* KO mice during social interactions**

(A, D) Exploratory pair-wise correlation matrix during nose-to-rear in WT mice (A) and *Mecp2* KO (E), nose-to-nose (B, F), over cagemate (C, G), and pushing behavior (D, H). Red arrows suggest possible clusters of neurons during each behavior. Pairwise correlations were performed using standard Pearson correlation between delta F/F traces.

**Supplemental Figure 6. Training protocol of the first cohort**

Differences was observed in trial duration and times of pushes during training between WT and KO mice, but not differences in the rate of change for both measurements. **(A)** Instrument and standard design of training in the tube test. **(B-C)** Trial duration during 3 days of training. **(C)** Trial duration grouped; Mecp2 KO mice spent significatively more time during trials than WT mice. **(D)** The rate of change in trial duration was similar between groups and within each group. **(E)** Times of pushes by trial during training **(F)** and grouped. **(G)** The rate of change of pushes was similar between groups and within each group. **(H)** Trial duration comparing groups and after ranking formation. Any difference by ranking during the daily trial duration and **(I)** grouped. **(J)** Pushes by group and after ranking formation. Any difference by ranking during the daily pushes and **(I)** grouped.

**Supplemental Figure 7. Training protocol of the second cohort**

Differences was observed in trial duration and times of crossing during habituation between genotypes WT and *Mecp2* KO mice, but not differences in the rate of change for both measurements. **(A)** Instrument and standard design of training in the tube test with small plastic cages at the ends of the tube. **(B-C)** Trial duration during 4 days of training. **(C)** Trial duration grouped; *Mecp2* KO mice spent significatively less time during trials than WT mice. **(D)** The rate of change in trial duration was similar between groups and within each group. **(E)** Times of crossing during habituation time; *Mecp2* KO mice crossing significatively more times during habituation than WT mice **(F)** and rate of change of crossing during habituation. **(G)** Trial duration comparing groups and after ranking formation. No one differences by ranking during the daily trial duration and **(H)** grouped.

