## Supplementary figures and images for "mPFC pyramidal neuron synchrony during social competition to form social rankings is disrupted in male *Mecp2* knockout mice"

### Supplemental Figure 1

#### Supplemental Figure 1. Male *Mecp2* KO mice have typical odor discrimination

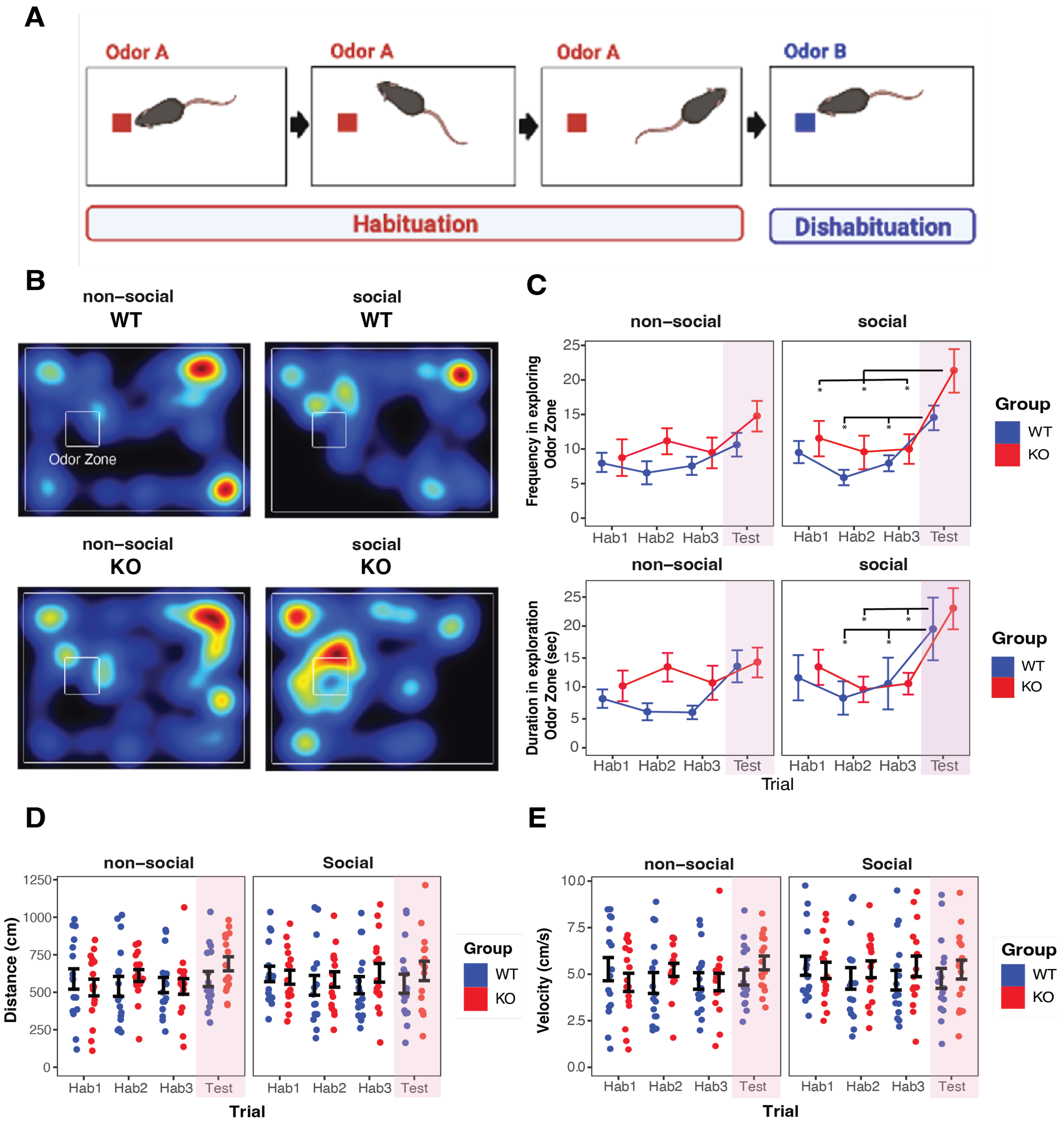

### Supplemental Figure 2

Supplemental Figure 2. Additional information on the warm spot test

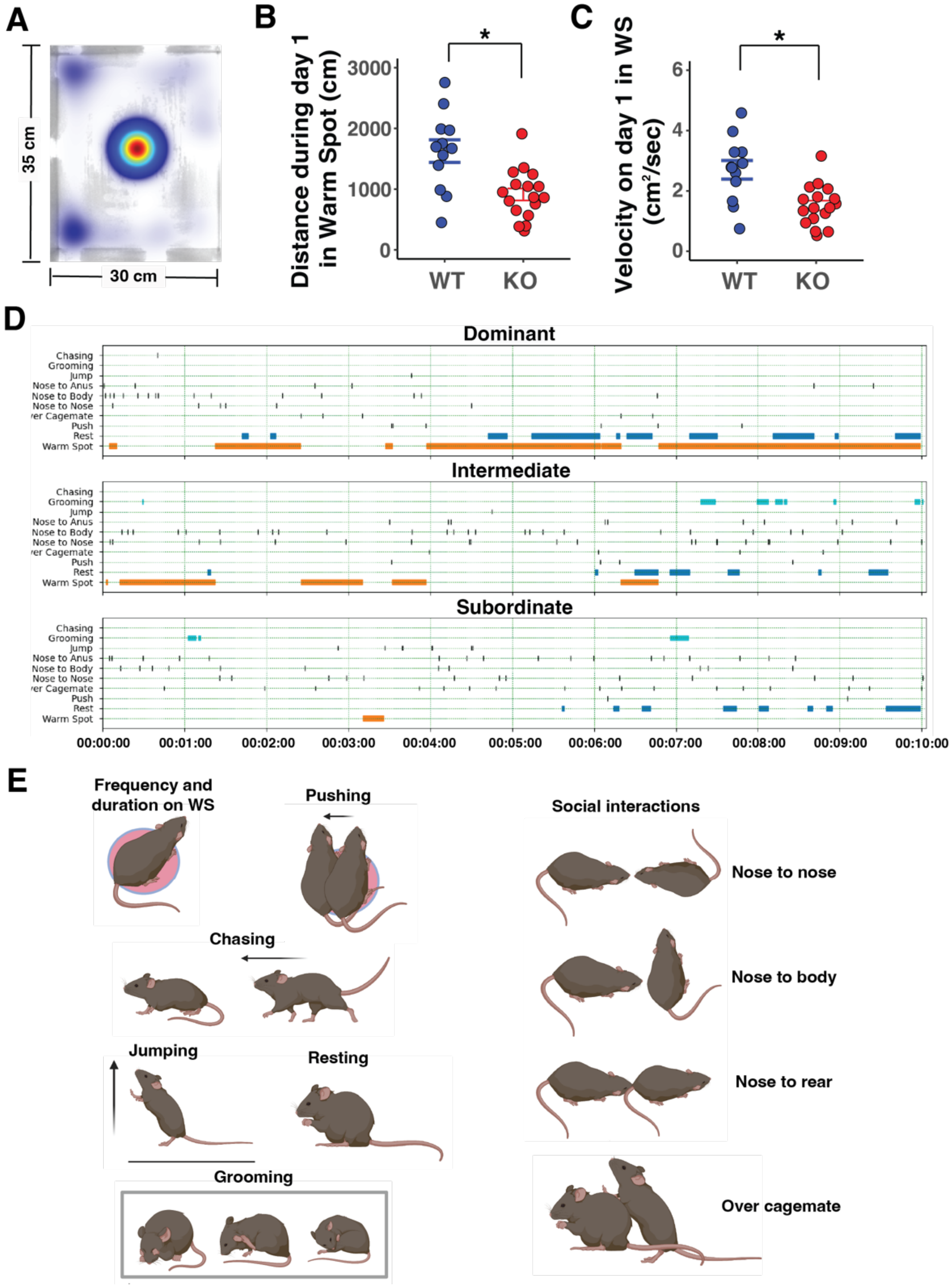

### Supplemental Figure 6

Supplemental Figure 6. Training protocol of the first cohort

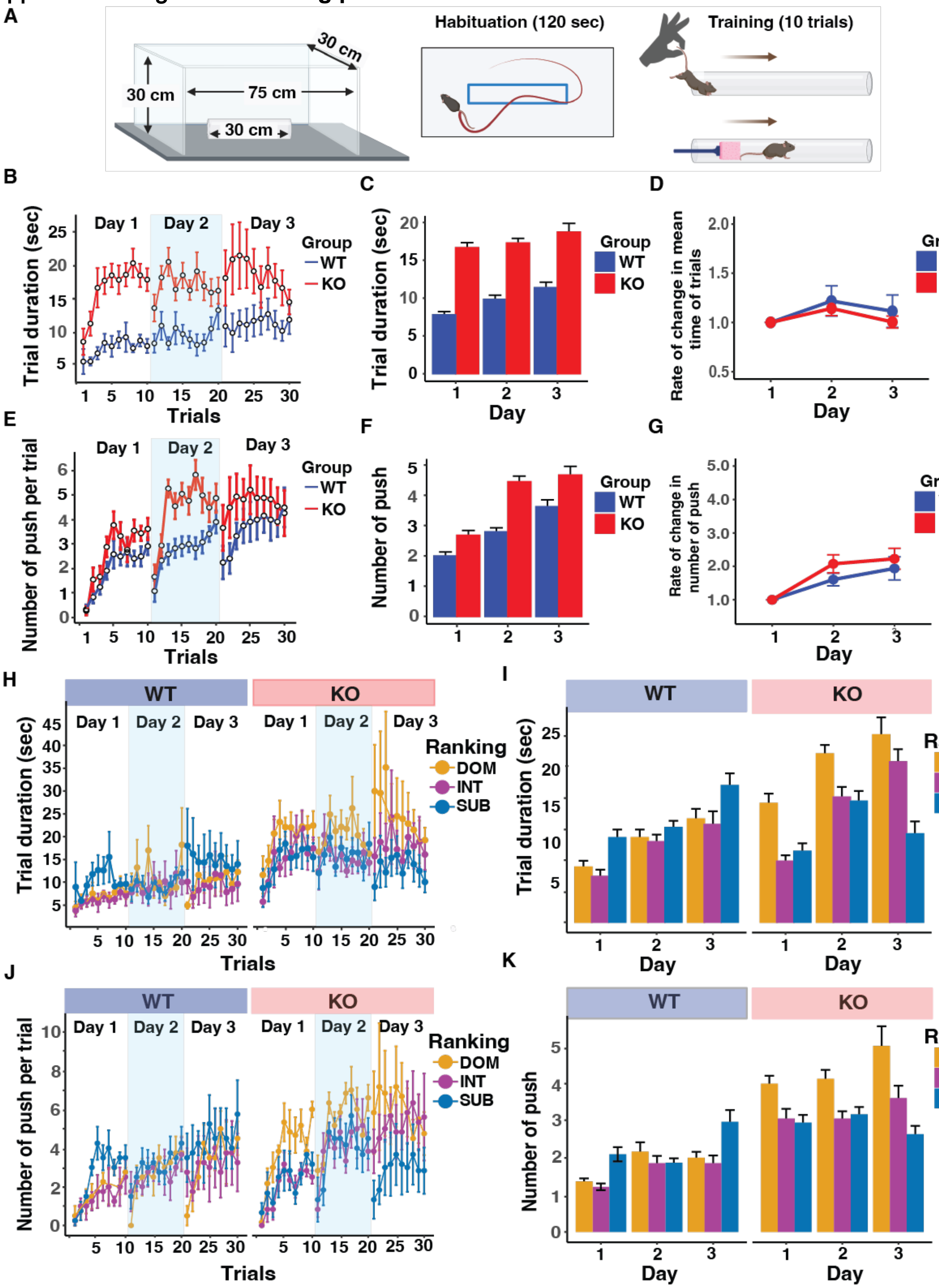

### Supplemental Figure 7

Supplemental Figure 7. Training protocol of second cohort

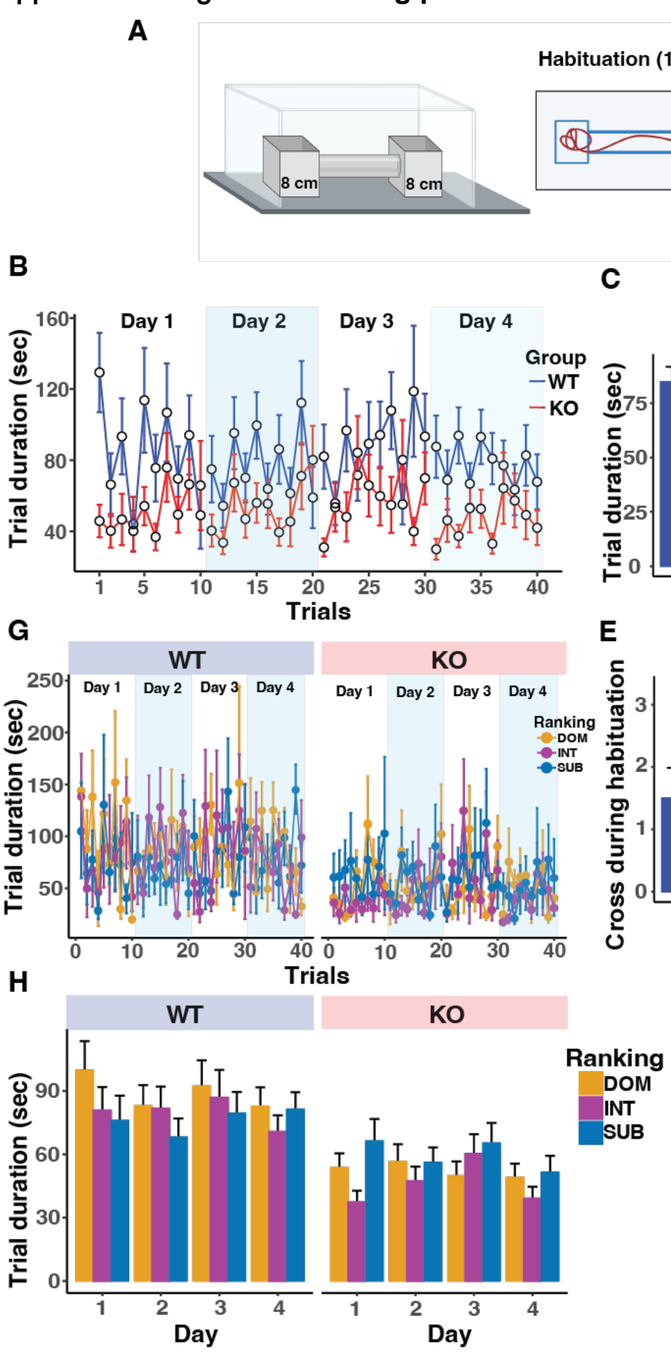
