## Supplemental Figure 3 for "mPFC pyramidal neuron synchrony during social competition to form social rankings is disrupted in male *Mecp2* knockout mice"

**Supplemental Figure 3. Socially sensitive neurons in mPFC active during social interactions were identified by *in vivo* $Ca^{2+}$ imaging with head-mounted miniscopes**

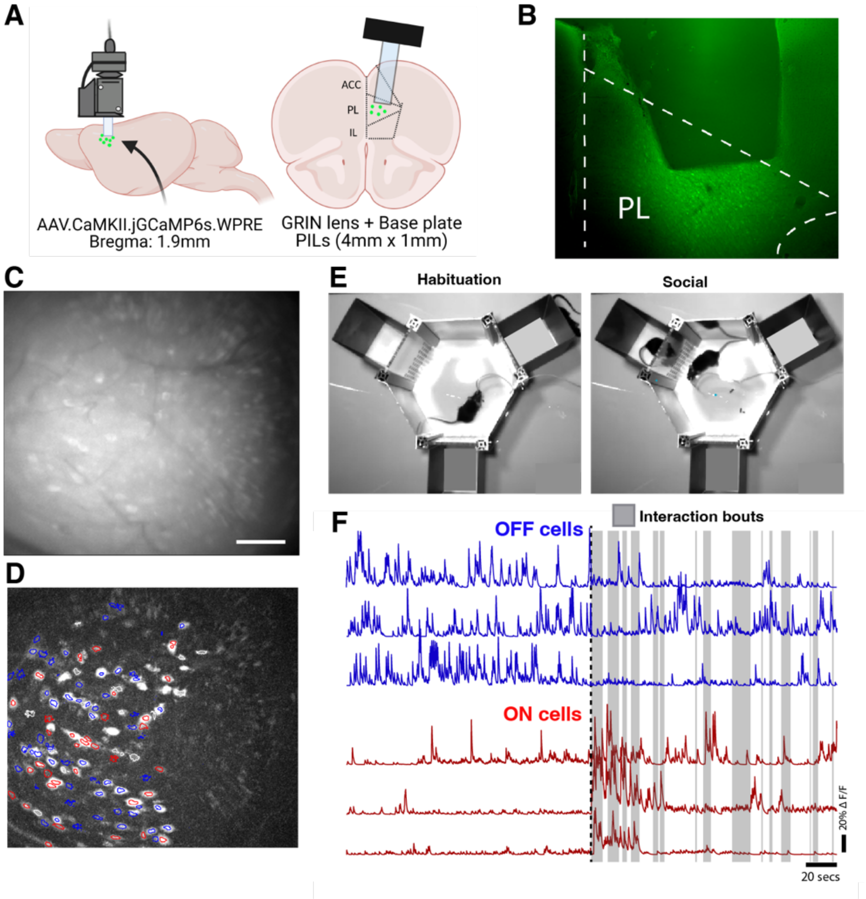
