## Supplemental Figure 4 for "mPFC pyramidal neuron synchrony during social competition to form social rankings is disrupted in male *Mecp2* knockout mice"

**Supplemental Figure 4. Social responsive neurons in mPFC were similar between *Mecp2* KO and WT mice, and similar dynamics between dominant and submissive social ranking**

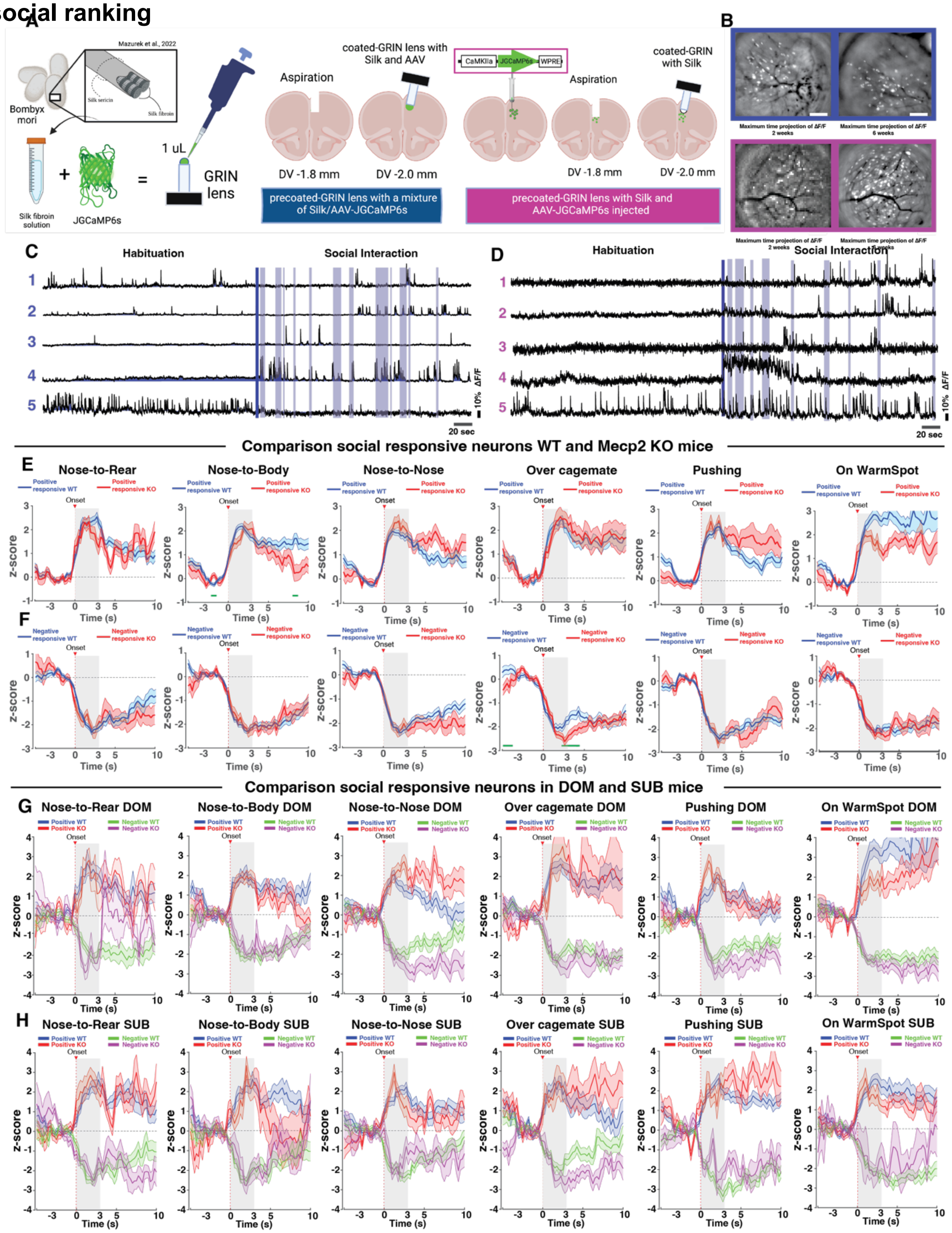
