## Supplemental Figure 5 for "mPFC pyramidal neuron synchrony during social competition to form social rankings is disrupted in male *Mecp2* knockout mice"

### Supplemental Figure 5. Pyramidal mPFC neurons showed higher correlation scores in WT mice than in *Mecp2* KO mice during social interactions

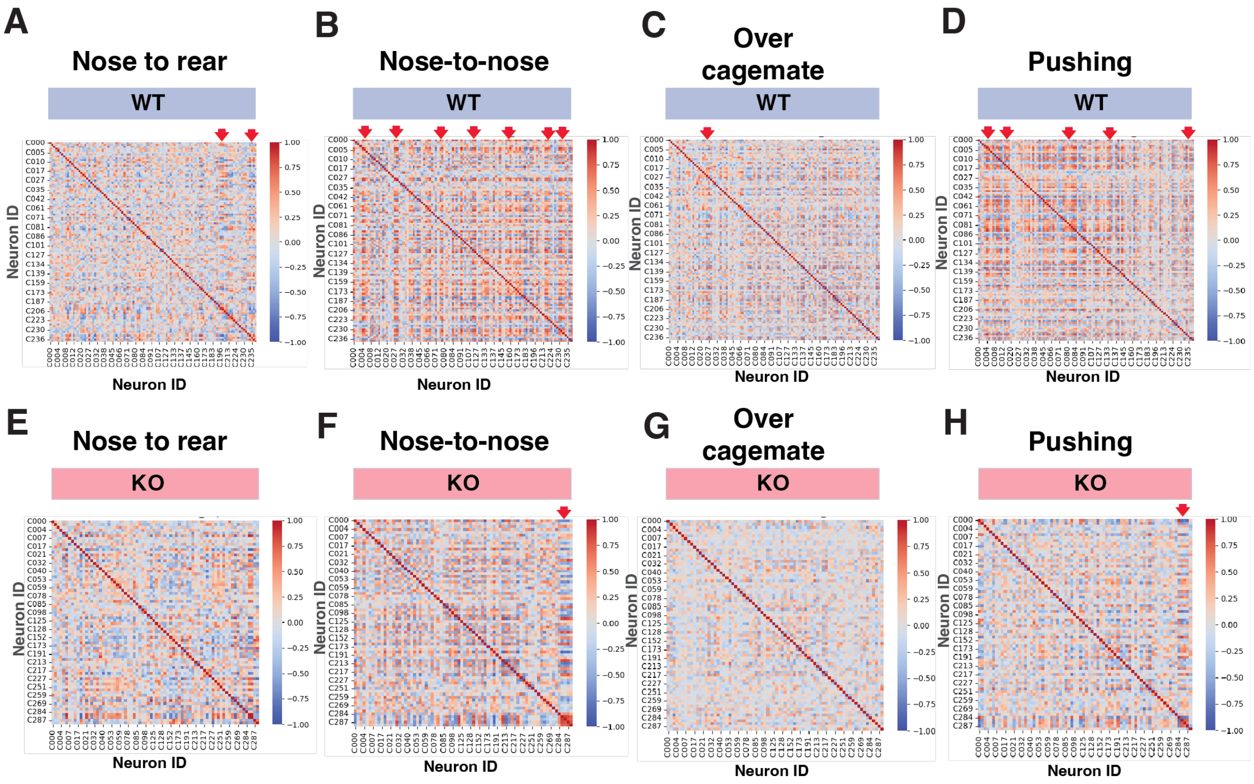
